# A tissue- and organ-based cell biological atlas of obesity-related human genes and cellular pathways

**DOI:** 10.1101/2020.03.16.993824

**Authors:** Iman Tavassoly, Valentina Barbieri, Coen van Hasselt, Pedro Martinez, Eric A. Sobie, Jens Hansen, Evren Azeloglu, Joseph Goldfarb, Philippe Sanseau, Deepak K Rajpal, Ravi Iyengar

## Abstract

Over the last decades, several features of obesity have been identified at behavioral, physiological, endocrine and genomic levels, and they have revealed the complexity of the disease; obesity results from a combination of genetic predisposition, endocrine disorders, and dysregulation of both food intake and energy expenditure. This complexity makes the development of new therapeutic regimens challenging and bariatric surgery is still the treatment of choice for many obese patients. Given the need for noninvasive therapeutic intervention strategies, we sought to systematically study the biological manifestations of obesity in peripheral organs. We analyzed publicly available datasets of genes, genomic determinants, and levels of obesity-related hormones in the blood, using a combination of methodologies, including graph theory and dynamical modeling, that allow for the integration of different types of datasets. The analysis revealed tissue- and organ-specific metabolic impairments and potential new drug targets. All the data are organized into a tissue/organ-based subcellular-function atlas for human obesity. The data show that the complexity of the obesity arises due to the multiplicity of subcellular processes in different peripheral organs.

## Introduction

Over the past decade, the prevalence of obesity and severe obesity among adults has risen to almost 40% of the world population [1]. According to the World Health Organization (WHO), obesity is a health issue not only in high income, but also in low- and mid-income countries, and is a risk factor for many chronic diseases including diabetes, cardiovascular diseases and cancer [2]. Obesity has been extensively studied at the behavioral, physiological, endocrine, and, more recently, the genomic level [3–6]. All these different biological levels and the interactions among them play a role in both the etiology and the pathophysiology of obesity and contribute to its complexity. Obesity is physiologically defined as a body mass index (BMI) of over 30kg/m^2^; it manifests itself as a combination of genetic predisposition, dysregulation of food intake and energy expenditure, and as an endocrine disorder making it challenging to develop therapeutic approaches to maintain body weight in a healthy range [3, 4]. After many years of trying, several large pharmaceutical companies have discontinued programs for the development of anti-obesity drugs [7]. Although there have been recent studies of peripheral drug targets for the treatment of obesity (incidental data have shown that GLP-1 agonists can be effective in moderate weight loss [8]), most FDA-approved drugs have primary targets in the Central Nervous System (CNS) [7]. Currently, the most effective treatment for obesity is bariatric surgery [9]. Given the need for new therapeutics that can control obesity without surgery, we hypothesized that we could identify potential novel therapeutic targets by systematically studying biological expressions of obesity in peripheral organs. We decided to utilize publicly available data because an enormous amount of information exists that can be analyzed using multiple computational systems biology approaches allowing for the integration of diverse types of datasets and models.

In principle, it is straightforward to argue that changes in genes and genomic determinants lead to alterations in cell biological pathways, and these, in turn, affect tissue and organ function, and finally whole-body physiology. However, it has been difficult, in practice, to connect the genomic information to the dynamics of physiological functions [10]. A recent review demonstrates the utility of integrated analyses of genomic determinants such as Single Nucleotide Polymorphisms (SNPs) with biological and behavioral determinants in obesity [11]. We sought to determine the relationships between genomic and molecular determinants and the dynamics associated with lean and obese physiological states. Data from patients after bariatric surgery provided a sample of subjects who had been obese but who had reduced their weight and BMI toward normal levels. As a measure of the physiological states, we used orexigenic and anorexigenic hormone levels in the blood and their changes after feeding. We hypothesized a) that genetic predisposition could be linked to endocrine impairment in energy homeostasis; b) that different tissues and organs could respond differently to genetic predisposition and endocrine impairment at the level of gene expression; c) that this variability could produce differences in catabolism and anabolism and confer distinct physiological profiles in lean, obese, and post-bariatric-surgery individuals. Because the effects of the same genetic and genomic determinants on cellular pathways may vary among different tissues and organs, we decided to map the genes and their associated cell biological processes to six peripheral tissues/organs that are known to be involved in nutrient processing and energy storage: stomach, intestine, pancreas, liver, adipose tissue, and skeletal muscle [12]. Upon performing this analysis, we organized the information as an atlas for human obesity, containing data from multiple biological levels.

The basic framework for the construction of the atlas is shown in **Figure 1**. The flow chart used for the analyses is shown in **Supplementary Figure S1,** and the databases used in the study are listed in **Supplementary Table S1**. The major goal of our mapping was to identify key genes and subcellular pathways by integrating the genomic and endocrine information using several computational methods. Genomic and genetic data were obtained from public databases [13–16] and then expanded utilizing the human interactome to take into consideration the role of possible intermediate genes. Because differences in endocrine regulation following food intake have been shown in lean, obese and post-bariatric surgery individuals [12], we determined whether such differences could be linked to genetic predisposition and how they would affect the genomic and genetic data. We used published data on changes in plasma levels of orexigenic and anorexigenic hormones to analyze physiological differences between lean and obese individuals and the effects of bariatric surgery on the plasma levels of hormones related to the regulation of food intake and obesity. We then developed a dynamical model that allowed us to identify those kinetic parameters that regulate the hormone levels in the systemic circulation and are more likely to be different in lean, obese, and post-bariatric surgery subjects. These findings allowed us to rank these kinetic parameters and the genes that are associated to them; the genes were then mapped onto the genomic and genetic network and weighted according to their significance in physiological functions; because we had three different physiological profiles, three endocrine-weighted genetic and genomic networks were obtained. To organize the information in terms of anatomy and biological functions, we filtered the ranked networks according to organ level transcriptomic patterns in humans [17]. We then used a combination of the FDA Adverse Events Database (FAERS) and DrugBank data [18, 19] to identify targets of drugs that have side effects of either weight gain or weight loss and drugs used for weight control whose targets include peripheral tissues and mapped these targets to the appropriate organs. The organ-selective gene lists were then used to identify relevant subcellular metabolic processes, leading to the creation of an atlas of obesity-related genes and subcellular processes in the stomach, pancreas, liver, adipose tissue, intestine, and muscle of the human body. The atlas illustrates how catabolic and anabolic subcellular processes differ at the tissue/organ level in lean, obese, and post-bariatric surgery individuals.

**Figure 1.**
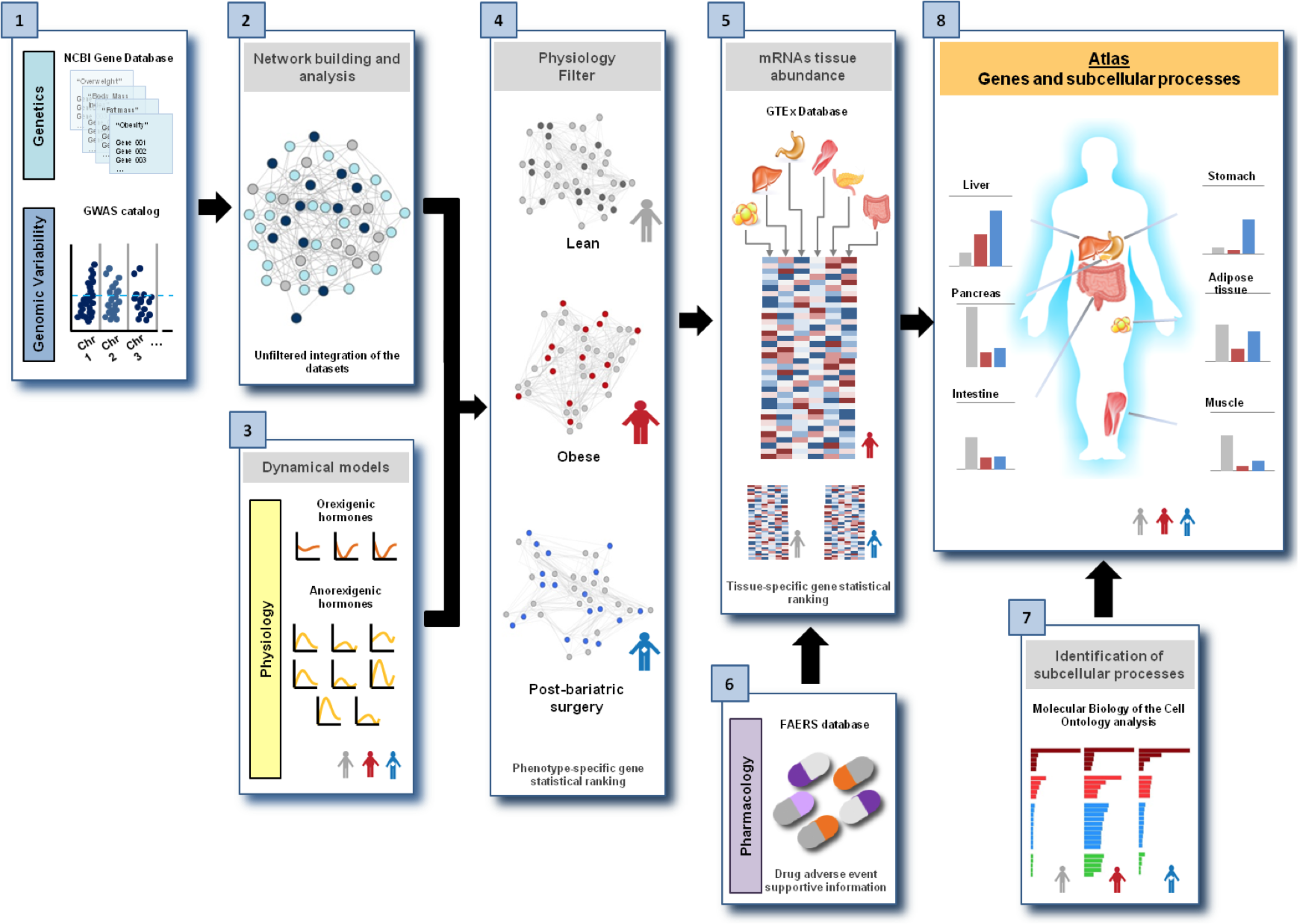
Schematic of the multi-model approach for mapping of genes and subcellular processes to different tissues and organs in the peripheral tissue atlas of metabolism. The atlas is generated through multiple types of data that are assembled in steps. **Step 1**: Integration of unweighted data sets. The genetic dataset (NCBI gene database) contains genes that are associated with leanness, obesity, overweight, body mass index, and fat mass. The genomic dataset (EBI GWAS) contains SNPs associated with obesity or with obesity-related traits with a *p*-value ≤ 5*10^-8^ according to published data. **Step 2:** Creation of unfiltered PPI network: The two datasets were statistically integrated by generating a genetic & genomic interaction network of the encoded proteins. **Step 3:** Generation of endocrinology data from modeling. The endocrinology dataset contains the genes associated with kinetic parameters for hormones that control satiety and hunger and that ranked differently in statistical significance in lean, obese, and post-bariatric surgery human subjects. These parameters were determined from an ODE-based multicompartmental dynamical model of the plasma concentrations of three orexigenic and eight anorexigenic hormones in lean, obese, and post-bariatric surgery human subjects. **Step 4:** Application of the physiology filter. The statistical significance ranking of genes in step 3 was used to weight each protein in the genetic & genomic network of step 2. Since the same gene can rank differently in lean, obese and post-surgery subjects, three subnetworks, one each for the lean, obese, and post-surgery phenotypes, were obtained. **Step 5:** Application of the tissue expression filter. The genes encoding each protein in each subnetwork were localized according to transcript abundance (GTEx database) in 6 organs/tissues: liver, pancreas, intestine, stomach, adipose tissue, and muscle. **Step 6**: Identification of known drug targets in the organ/tissue-specific subnetworks: A dataset extracted from FAERS database of molecular targets of drugs affecting body weight was used to support the analysis from the previous steps. **Step 7:** Application of the subcellular processes filter. We conducted enrichment analysis using Molecular Biology of the Cell Ontology (MBCO) to identify the subcellular processes that are significantly associated with a lean, obese, and post-surgery phenotype in each organ/tissue. **Step 8:** Generation of the peripheral obesity atlas. Integration of these data enable the application of statistical models to generate graphical representations of differences in genes associated with different organs/tissues in the lean, obese, and post-surgical condition, which could be displayed on a peripheral obesity atlas.

## Results

### Genetic and genomic datasets

We searched the NCBI Gene database and the EBI GWAS catalog (**Supplementary Table S1**) [13–16] to identify genetic and genomic variability associated with BMI in humans. The NCBI Gene database contains genes that have been associated with specific traits in published studies (2006-2018). The GWAS catalog includes all the SNPs and mapped genes associated with diseases or traits in published studies from 2006 to 2018. The overall strategy for identifying relevant genes is shown in **Supplementary Figure S2**. As of April 2018, we identified 2527 genes associated with either “obesity”, “body mass index”, “overweight”, “free fatty mass”, or “leanness” from the NCBI Gene database (**Supplementary Figure S2**). The complete list of query terms and the number of associated genes is shown in **Supplementary Table S2**. The list of genes and associated publications are shown in **Supplementary File 1**. Genomic variability was incorporated as SNPs associated with obesity, leanness, and obesity-related traits with a *p*-value ≤ 5*10^-8^ from the EBI GWAS catalog (**Supplementary Figure S3**). The SNPs were mapped to a total of 324 regions that were utilized for subsequent analysis (**Supplementary Figure S3, Supplementary Table S3, Supplementary File 2**). Common to both sets were 117 genes, yielding 2734 unique genes (**Figure 2A**) that are distributed among all chromosomes except the sex chromosomes (**Figure 2B)**. To further identify all the genes related to pathways involved in obesity, we built a protein-protein interaction (PPI) network, using proteins encoded by the 2734 unique genes from the two datasets as seed nodes. Using the nearest neighbor expansion method, we then identified proteins in the human interactome that were one intermediate node between seed nodes. This process resulted in a network of 7268 gene products that, at this stage of the study, were considered to have equal statistical significance and constituted the main dataset for subsequent analyses (**Figure 2C and Supplementary Figure S4**). To identify the major biological processes that these genes participate in, we performed an enrichment analysis for biological processes utilizing the Molecular Biology of the Cell Ontology (MBCO) algorithm [20]. This algorithm allows for the identification of interactions between subcellular processes (SCPs) and classifies them into hierarchical levels according to their function. The analysis showed that the genes from the unfiltered PPI network dataset seemed to be significantly related to cellular communication, lipid metabolism, cellular response to stress, extracellular matrix homeostasis, metabolism, and transport of glucose and lipids (**Figure 2D and Supplementary Figures S5, S6, and S7)**.

**Figure 2.**
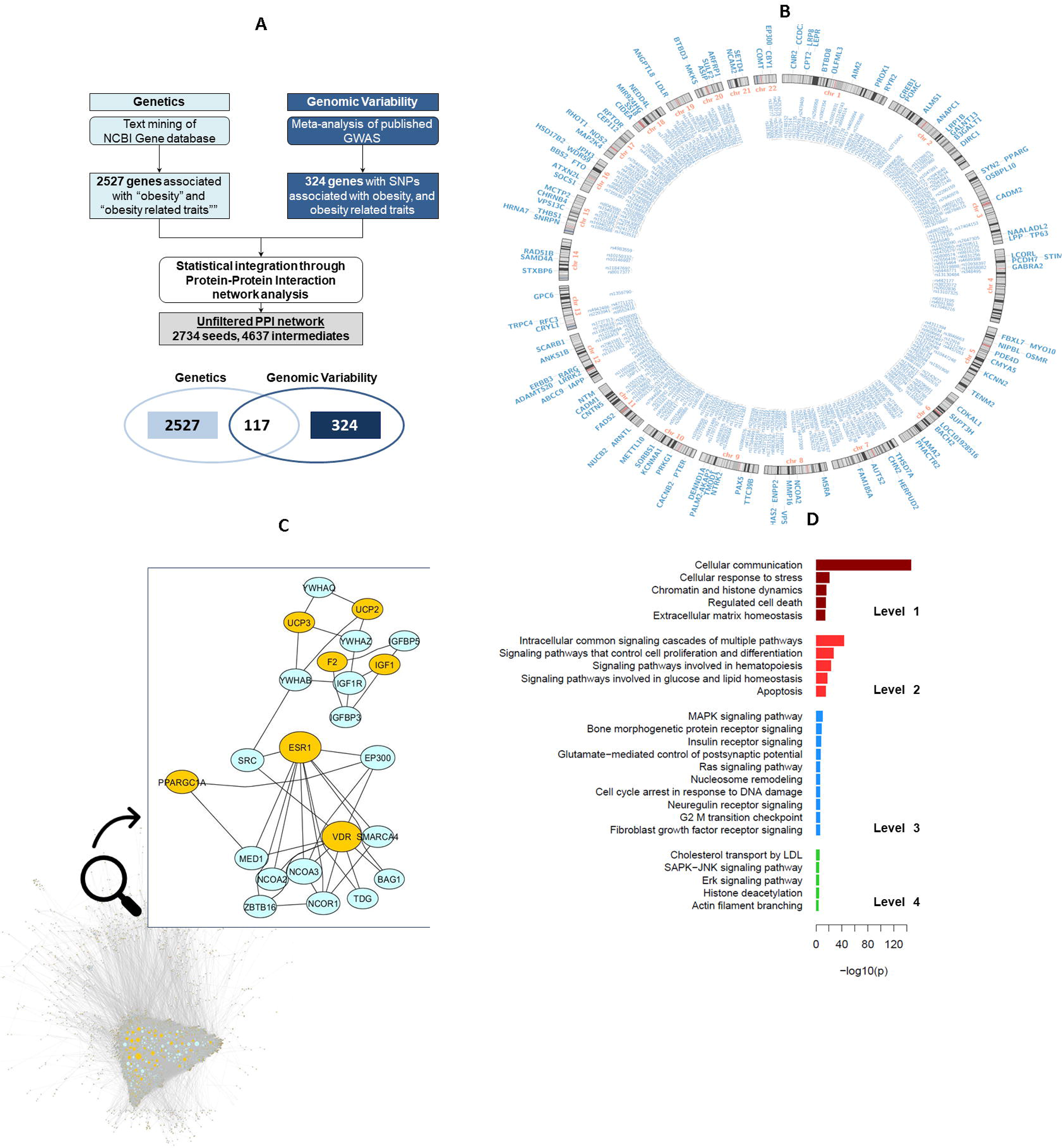
Survey of genes and genomic variants associated with human obesity and weight - related traits. **A) Flowchart of the information process.** We identified 2527 genes associated with “obesity” (BMI ≥ 30kg/m^2^), “body mass index,” “overweight,” “free fatty mass,” or “leanness” (BMI ≤ 25 kg/m^2^), in at least two publications from the NCBI Gene database (https://www.ncbi.nlm.nih.gov/gene). Genomic variability was incorporated as SNPs associated with obesity or obesity-related traits with a *p*-value ≤ 5*10^-8^ from the EBI GWAS catalog (http://www.ebi.ac.uk/gwas/). Mapping the SNPs to obesity and obesity-related traits identified 324 genes, 207 of which were not in the original NCBI set. The two datasets were integrated through construction of a PPI, using the nearest neighbor expansion method with the proteins encoded by the genes as seeds and identifying intermediate proteins (and their encoding genes) from the human interactome up to two path lengths of distance. This process resulted in a network of 7268 proteins (and encoding genes); seed genes that lacked connection to the network were not used for further analysis. **B) Mapping the PPI network onto chromosomes.** The outermost layer contains the gene symbols (blue text) of genes associated with obesity and/or obesity-related traits according to the NCBI Gene database. The next layer shows the chromosome bands (colored rectangles) of 22 human chromosomes. Chromosome 23 and the Y chromosome do not contain genes of interest and were omitted. The next layer lists the SNPs associated with obesity (Blue). The final innermost layer lists the SNPs associated with obesity-related traits. SNPs were assigned according to the GWAS catalog. **C) Unfiltered PPI network.** Representation of the unfiltered PPI network: Seed nodes from the GWAS and NCBI datasets are in yellow, intermediate genes are in light blue. **D)** Enrichment analysis of the unfiltered PPI network for subcellular processes. The MBCO algorithm was used to identify which subcellular processes the gene products in the unfiltered network belong to. The algorithm screens the PubMed titles and abstracts of each SCP-specific article set and counts articles mentioning a given gene at least once. The number of abstracts needed to associate a gene with a process varies with the level of the SCP. A minimum of 4 abstracts is required for a level-1 SCP, 3 abstracts for a level-2 SCP, 2 for a level-3 SCP and 1 for a level-4 SCP. Fisher’s exact test is used to calculate *p*-values.

### Endocrine dynamics datasets

Impairment in hormonal regulation of hunger and satiation is a pivotal physiological marker of obesity [12]. One reasonable explanation for the observed differences between lean, obese, and post-bariatric surgery states is that kinetic regulation of hormones could be different in these three physiological states. To assess this, we developed a dynamical model that describes the basal secretion, stimulus-evoked secretion, utilization, and degradation of 3 orexigenic and 8 anorexigenic peripheral hormones in human subjects [8, 21–32]. Synthesis and storage of the hormones were not considered explicitly but were subsumed as part of the secretion rate. Our goal in developing this dynamical model was to identify the kinetic parameters to which hormone levels were most sensitive. These kinetic parameters, in turn, can be associated with cellular pathways and genes related to obesity. Our general strategy was to choose parameter values in the model by fitting model time courses to experimentally measured blood hormone levels, before and after a meal. As we do not know if these are the true biologically relevant parameters, we then conducted extensive parameter variations followed by multivariable regression analyses to identify the most sensitive kinetic parameters associated with the levels of each hormone.

We collated plasma level concentrations at seven time points from published studies [8, 24–32] to include a fasting state (−120min-0min), a feeding point (at 0 min) and a post-prandial state (0-120min). Four hours is the usual timeframe for blood sample collection in published studies, so we ran our simulations accordingly. The model reproduces the biological response to the ingestion of glucose as follows: glucose stimulates ghrelin (GHRL) secretion from the stomach. Absorption by the intestine stimulates the secretion of glucagon-like peptide-1 (GLP-1), cholecystokinin (CCK), oxyntomodulin (OXM), gastric inhibitor peptide (GIP), and peptide YY (PYY). Once absorbed, glucose delivery to the pancreas via the mesenteric circulation stimulates the secretion of insulin (INS), glucagon (GCG), and amylin (AMY), and delivery to adipocytes via the systemic circulation stimulates secretion of leptin (LEP) and adiponectin (ADIPOQ) (**Figure 3A**). According to the literature, in the lean subjects, orexigenic hormones (GHRL, ADIPOQ, and GCG) show higher fasting levels and lower post-prandial levels. The anorexigenic hormones (INS, GLP-1, CCK, OXM, GIP, PYY, and LEP) present lower fasting levels and higher postprandial levels. In obese subjects, hormone responsiveness to nutrients was reported to be totally (GHRL, ADIPOQ, GLP-1, PYY, OXM) or partially impaired (GCG, LEP, INS, AMY, CCK, GIP). Plasma levels of some hormones in post-bariatric surgery patients lie between the lean-curves and the obese-curves (GHRL, GCG, LEP, AMY, GIP), while others appear higher than in either lean or obese individuals (INS, CCK, GLP-1, PYY, and OXM) (**Figure 3B**). We ran simulated time courses for lean, obese and post-bariatric surgery subjects in a fasting and a postprandial state to estimate reliable initial values for the kinetic parameters such that the simulated curves closely resembled published experimental data [8, 25, 27, 28, 30–33] **(Figure 3B and Supplementary File 3**).

**Figure 3.**
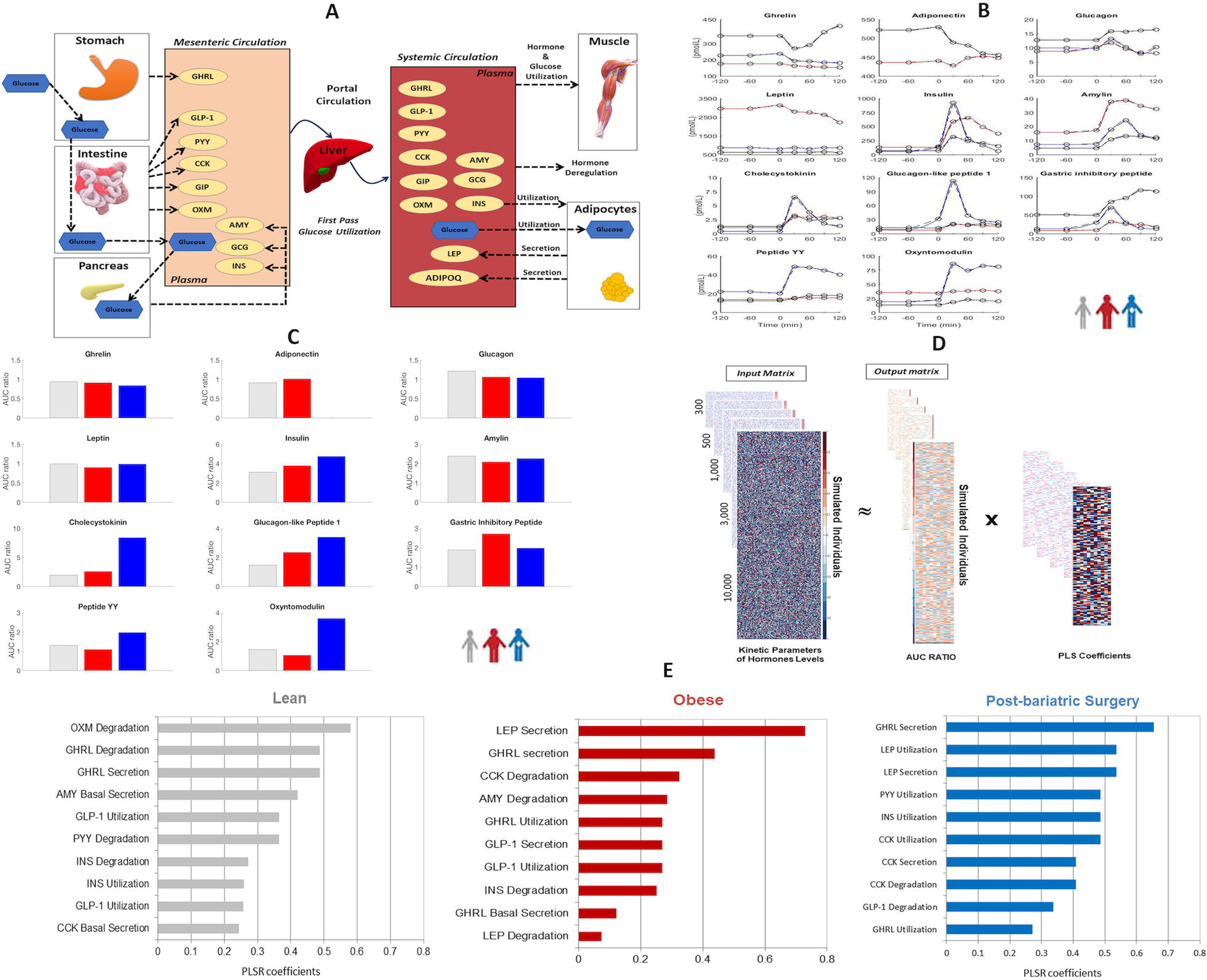
Dynamical modeling of temporal patterns of plasma concentrations of hormones associated with food intake in lean, obese and post-bariatric surgery patients. **A) Compartmental model of the plasma levels of selected peripheral hormones in human subjects**. Glucose-responsive peripheral hormones that control food intake are modeled with the kinetic parameters of secretion, utilization, and degradation. Ingestion of glucose stimulates ghrelin (GHRL) secretion from the stomach. Absorption by the intestine stimulates the secretion of glucagon-like peptide-1 (GLP-1), cholecystokinin (CCK), oxyntomodulin (OXM), gastric inhibitor peptide (GIP), and peptide YY (PYY). Through mesenteric blood circulation, glucose delivery to the pancreas stimulates the secretion of insulin (INS), glucagon (GCG), and amylin (AMY) and delivery to adipocytes stimulates secretion of leptin (LEP) and adiponectin (ADIPOQ). All the hormones, except for leptin and adiponectin are subjected to first pass through the liver, and then they are delivered by the circulation to the muscles for utilization, or they undergo degradation. (No data was available of measuring ADIPOQ in the post-bariatric surgery subjects) **B) Simulated time courses of the plasma concentrations of the hormones**. Simulated time courses for lean (black with gray circles), obese (red) and post-bariatric surgery (blue) subjects are shown. Simulations were conducted using parameters chosen so that the curves closely resembled experimental data (in dotted lines) from different studies (**Supplementary File 3**). Subjects are in a fasting state from −120 to +0 min. At 0 minutes, administration of glucose initiates the postprandial state, which is monitored for 120 min. **C) Quantitative output of the simulation of the dynamical model.** Ratio of the area under the curve (AUC) of the postprandial plasma hormone concentration (2 hours) divided by the AUC of the fasting plasma concentration (2 hours for each hormone was chosen as the quantitative output of the simulation for use in the multivariable regression analysis (MRA). **D) Multivariable regression analysis (MRA) of simulated data identifies kinetic parameters whose variation produces the greatest changes in glucose-evoked hormone release in lean, obese, and post-bariatric surgery subjects.** To account for the variation in the kinetic regulation of hormone physiology in a population, and to compensate for the lack of extensive clinical data, we generated 3 simulated sets of human subjects (lean, obese, post-surgery) by randomly varying each of the 141 kinetic parameters of the model, with the mean centered on the parameter value and the standard deviation set to ± 1. We used partial least square regression (PLSR) to identify which kinetic parameters are more likely to induce a significant perturbation of the output of the dynamical model (AUC ratio). MRA was repeated with progressively larger populations (representing 300, 500, 1,000, 3,000, 10,000 individuals). The MRA for the obese population of 10,000 is represented here: We generated a total of 10,000 parameter sets that were stored in the input matrix, *I(10,000×141)*. Each new set of parameters was then utilized to simulate the dynamical model, and the output (AUC ratios for each hormone) was stored in the output matrix, *O(10,000×11)*. The PLSR, performed on matrices *I* and *O*, produced a matrix of regression coefficients, *B_PLS_ (141×11)*, that can be used to predict the resulting outputs such that *O_predicted_* = *I x B_PLS_* is close to the original output matrix *O*. **E) MRA ranking in each subject population.** The top 10 most sensitive kinetic parameters from simulations of lean (grey), obese (red), and post-bariatric surgery (blue) subjects, according to PLSR coefficients determined through multivariable regression analysis of the ratio between the area under the curve (AUC) of the postprandial plasma hormone concentration and the AUC of the fasting plasma hormone concentration. The resulting PLSR coefficients were then used to rank the parameters.

The magnitude of the endocrine response to nutrients can be assessed by the difference in plasma levels of the hormones before and after the meal. To quantify it, we chose to calculate the ratio between the 2-hour area under the curve (AUC) of the postprandial plasma concentration and the 2-hour AUC of the fasting plasma concentration for each hormone (**Figure 3C and Supplementary Figure S8)**. To identify which kinetic parameters are more likely to affect the endocrine response represented by the AUC, we performed a multivariable regression analysis (MRA) [34]. The logic of this analysis was to randomly vary the initial values of the set of kinetic parameters used in the model a number of times (**Supplementary Figure S9**), utilizing each new set of values to run a simulation that by varying each parameter affects the output and expresses it in terms of a numerical Partial Least Square Regression (PLSR) coefficient: the higher the coefficient, the more sensitive is the response to the parameter; the lower the coefficient, the more robust is the response to changes in the parameter. The kinetic parameters for lean, obese, and post-surgery populations were then ranked according to the coefficients. The methodology is schematically shown in **Figure 3D and Supplementary Figure S10** and fully explained in online methods.

We focused on the top ten most sensitive kinetic parameters from simulations of lean, obese, and post-bariatric surgery subjects because they account for 90% of the total sensitivity **(Figure 3E)**. This analysis can indicate both the processes that have the greatest effects on hormone levels and how their relative importance can be different between physiological states. For instance, most of the top-ranking parameters involved the degradation and utilization of anorexigenic hormones, except for ghrelin, which ranked highly in all three physiological states. In lean subjects specifically, oxyntomodulin and amylin also seem to play an important role, while insulin ranked lower. In obese subjects, leptin secretion scored the highest, but cholecystokinin and amylin were among the top five. Glucagon-like peptide 1 seems to play a key role as well, providing an unbiased physiological reason for its therapeutic use for obesity [35]. In post-surgery subjects, utilization of hormones secreted by the intestine appeared to play a key role, and it could, in part, be related to the changes in the anatomy of the patients due to the surgery.

### Mapping of the physiological responses onto the genetic and genomic determinants

We determined whether there were correlations between the genetic and genomic dataset (**Figure 2C**) and the physiological regulation results (**Figure 3E**). To identify genes that are associated in the literature with each kinetic parameter, we text-mined the PubMed database utilizing the MBCO algorithm [20]. For each parameter, we obtained a list of genes ranked by their association to it based on published data (**Figure 4A**). Each gene was then assigned an average rank defined as the mean of the MBCO rank and the MRA rank for the associated parameter (**Figure 4A**). Because the same gene can have a different average rank in lean, obese, and post-surgery subjects due to different MRA ranking of the parameter in the three populations, we obtained population-specific gene ranks that incorporated their predicted endocrine sensitivity (**Figure 4B**). The complete list of genes and parameters is in **Supplementary File 4.**

**Figure 4.**
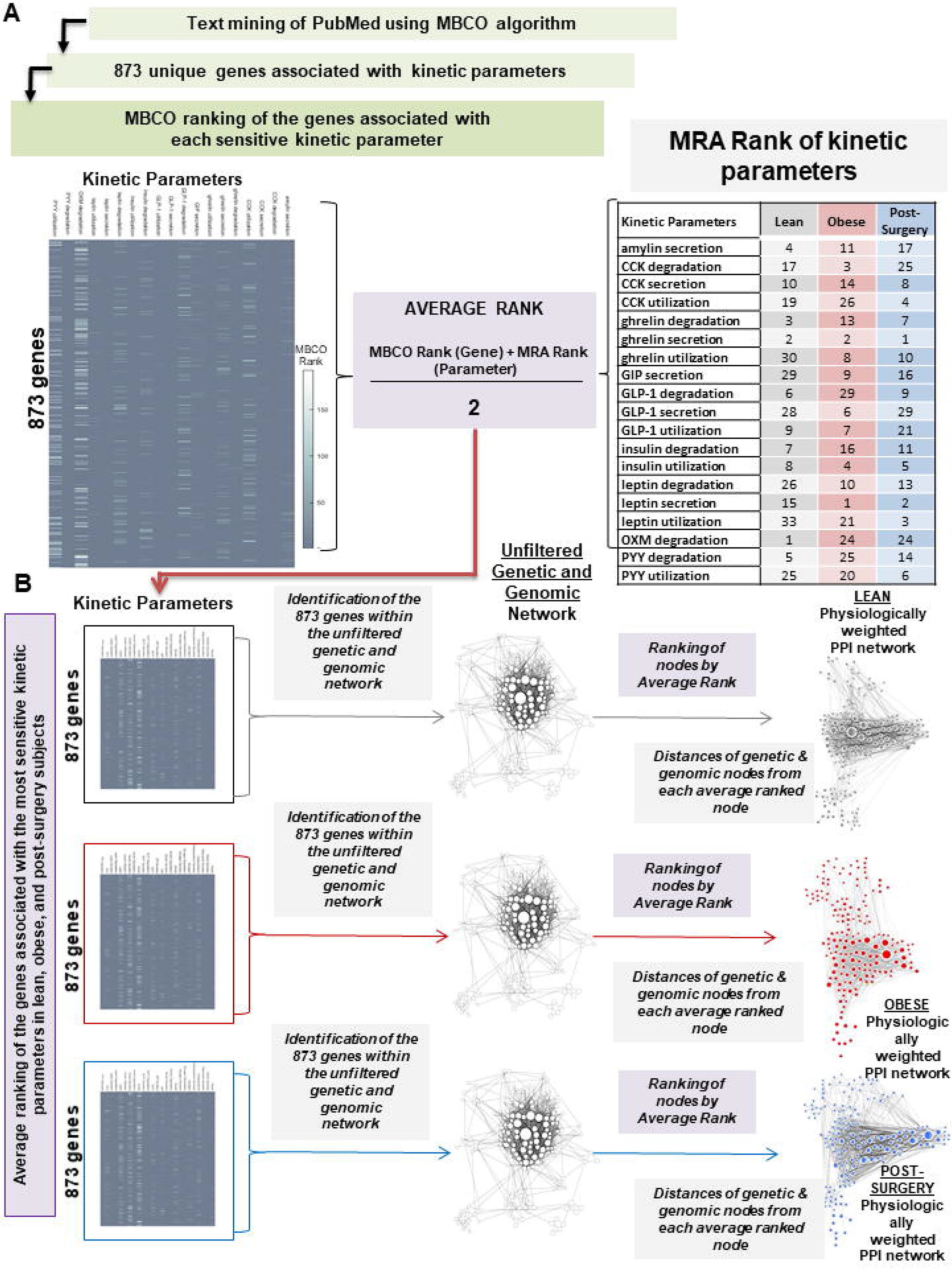
Integration of the PPI network and the dynamical model data to obtain physiologically weighted tissue-specific PPI networks for lean, obese, and post-bariatric surgery subjects. **A) Average ranking of genes associated with the most sensitive kinetic parameters in lean, obese, and post-surgery subjects. Top left.** The genes associated with each of the sensitive parameters were obtained utilizing the Molecular Biology of the Cell Ontology (MBCO) algorithm and were ranked according to their relative MBCO score. **Top right.** The most sensitive kinetic parameters in lean, obese, and post-surgery subjects were ranked according to their PLSR rank obtained from the MRA. **Top center.** The average ranking score for each gene was calculated as the mean of the MBCO rank and the MRA rank. **B) Weighting of the encoding genes in the unweighted PPI network using the results from dynamical modeling analysis and MBCO classification. Bottom left:** The average rank score for each gene (left) was used to weight each corresponding node in the unfiltered genetic & genomic PPI network (**center**). Since the same gene can rank differently in lean, obese, and postsurgery subjects, the analysis is performed separately for each condition. The remaining nodes encoding proteins within the lean-, obese-, and post-surgery PPI networks were assigned a rank according to their distances from each of the average ranked genes *(edge thickness)* Distances are calculated from 1 to *n* and generate a physiologically weighted PPI network for each physiological condition (lean, obese, post-surgery) (**right**).

We looked for the genes identified in the endocrine sensitivity analysis in the unfiltered genetic and genomic PPI network **(Figure 4B)** and weighted them using the average rank previously calculated. Because these rank values are population-dependent, we generated lean-, obese-, and post-surgery PPI networks (**Figure 4B**). The genes that are present in the endocrine dataset but not in the genomic and genetic dataset were weighted according to their distance from the ranked genes. This mapping allowed us to classify the genetic and the genomic dataset according to its connection with the endocrine ranked genes and to create population-specific networks: genes that are present in both datasets ranked at the top; genes that are present in the endocrine dataset but not in the genomic and genetic dataset ranked lower.

Because the same gene products can differ in expression, regulation, and function in different tissues which have significant roles in weight control, we investigated which genes in the 3 PPI networks were expressed in peripheral tissues/organs of interest. We obtained the tissue/organ-specific mRNA expression data for each gene from the GTEX portal for adipose tissue, intestine, liver, skeletal muscle, pancreas, and stomach (**Supplementary Figure S11**). We divided the genes of the three subnetworks according to their expression in the 6 tissues/organs. The analysis generated 18 PPI subnetworks, one for lean, obese, and post-surgical subjects in each of the 6 tissues.

While most weight-loss drugs that have been marketed have primary targets in the CNS, it may be advantageous to seek peripherally active therapeutic approaches. We determined whether any FDA-approved drugs that have a high risk of weight gain or weight loss as side effects have as secondary targets, gene products in any of the subnetworks (**Supplementary File 5)**. We selected drugs that have exclusively peripheral targets and drugs that have both peripheral and Central Nervous System (CNS) targets. Drugs with exclusively CNS targets, as lorcaserin and phentermine, were excluded. The respective targets for each drug were obtained from Drugbank [19] and DGIdb [36]. The top 10 drugs associated with weight gain and weight loss, and their targets are presented in **Figure 6**. We mapped the targets onto the tissue-specific subnetworks (**Supplementary Figure S12 and S13**). 8 weight gain-associated targets and 11 weight loss-associated targets are distributed among the 6 tissues/organs of interest so that each contains at least one target **(Figure 5)**. The implications of this result are two-fold: first, the presence of known secondary targets in the subnetworks generated by our analysis indicates that one reason behind the complex pathophysiology of obesity may be that determinants can be identified in several levels. Second, because the CNS-associated side effects of anti-obesity drugs are undesirable, it suggests the possibility that we can identify new peripheral targets to treat obesity.

**Figure 5.**
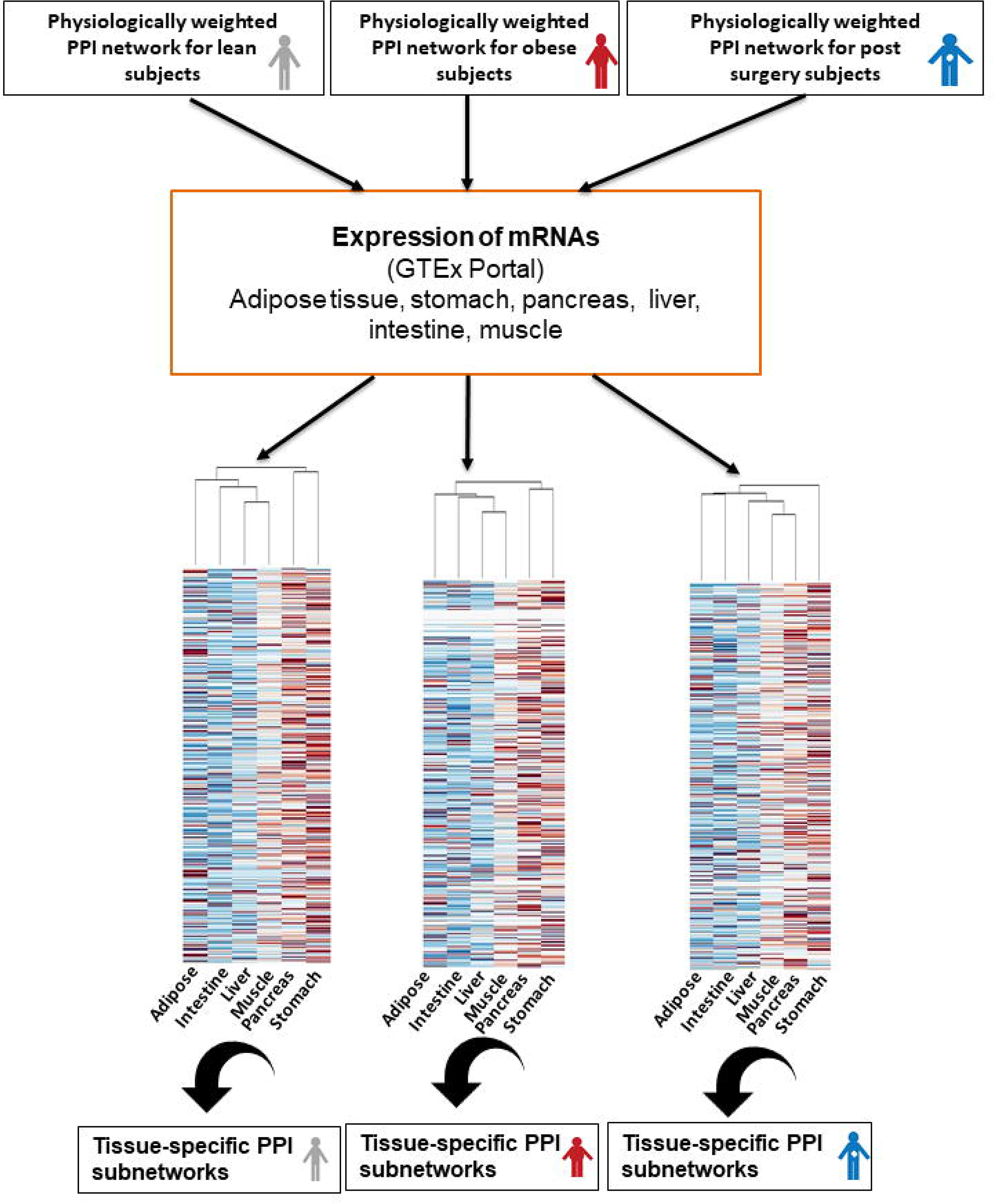
Organ/tissue-specific gene expression patterns in the physiologically weighted PPI networks. The physiologically weighted PPIs for lean, obese, and post-surgery conditions were divided according to the expression of each gene in the indicated tissues/organs, resulting in the generation of a PPI subnetwork for each site.

### Catabolic and anabolic differences in lean, obese and post-surgery subjects

To evaluate differences in catabolism and anabolism in our PPI models of lean, obese and postsurgery subjects, we identified, utilizing the MBCO algorithm, the subcellular processes that the gene products in the 18 subnetworks participate in [20] (**Supplementary Figure S14**). We classified these metabolic processes as catabolic and anabolic processes, according to standard textbook criteria [37]. Ratios of catabolic and anabolic processes within the top 100 subcellular processes within each of the six tissues and organs of interest and each physiological condition were calculated.

Interestingly, lean and post-surgery subjects show a higher number of subcellular catabolic processes, while obese subjects have similar numbers of each (**Figure 7**). When the subcellular processes are analyzed at the organ level, results vary considerably, and the apparent similarity of lean and post-surgery subjects is lost. In lean subjects, catabolic processes seem to be favored in pancreas, intestine, and muscle, but with ratios close to 1 in liver, stomach and adipose tissue. In post-surgery subjects, catabolic processes predominate in liver, stomach, and muscle, ratios are close to 1 in the intestine and adipose tissue, and anabolic processes predominate in the pancreas. These differing profiles merit further study. In obese subjects, catabolic processes predominate only in the liver, anabolic in stomach, pancreas, and adipose tissue, and ratios approach 1 in muscle and intestine. The large change in the catabolic/anabolic ratio in the stomach in post-surgery versus obese or lean subjects could reflect changes in the gastrointestinal anatomy due to surgical removal of a portion of the stomach. Our results suggest that anabolic and catabolic processes in the pancreas, intestine, and muscle could be considered as potential peripheral targets for future combination therapeutic treatments, since in obese subjects the catabolic/anabolic ratio in these organs is less than 1 meaning the anabolism is dominant while in the lean subjects this ratio is higher than 1 in these organs meaning that catabolism is dominant. These organs can be targeted to change the ratio and balance between the catabolism and anabolism in obese subjects. (**Figure 7**)

**Figure 6:**
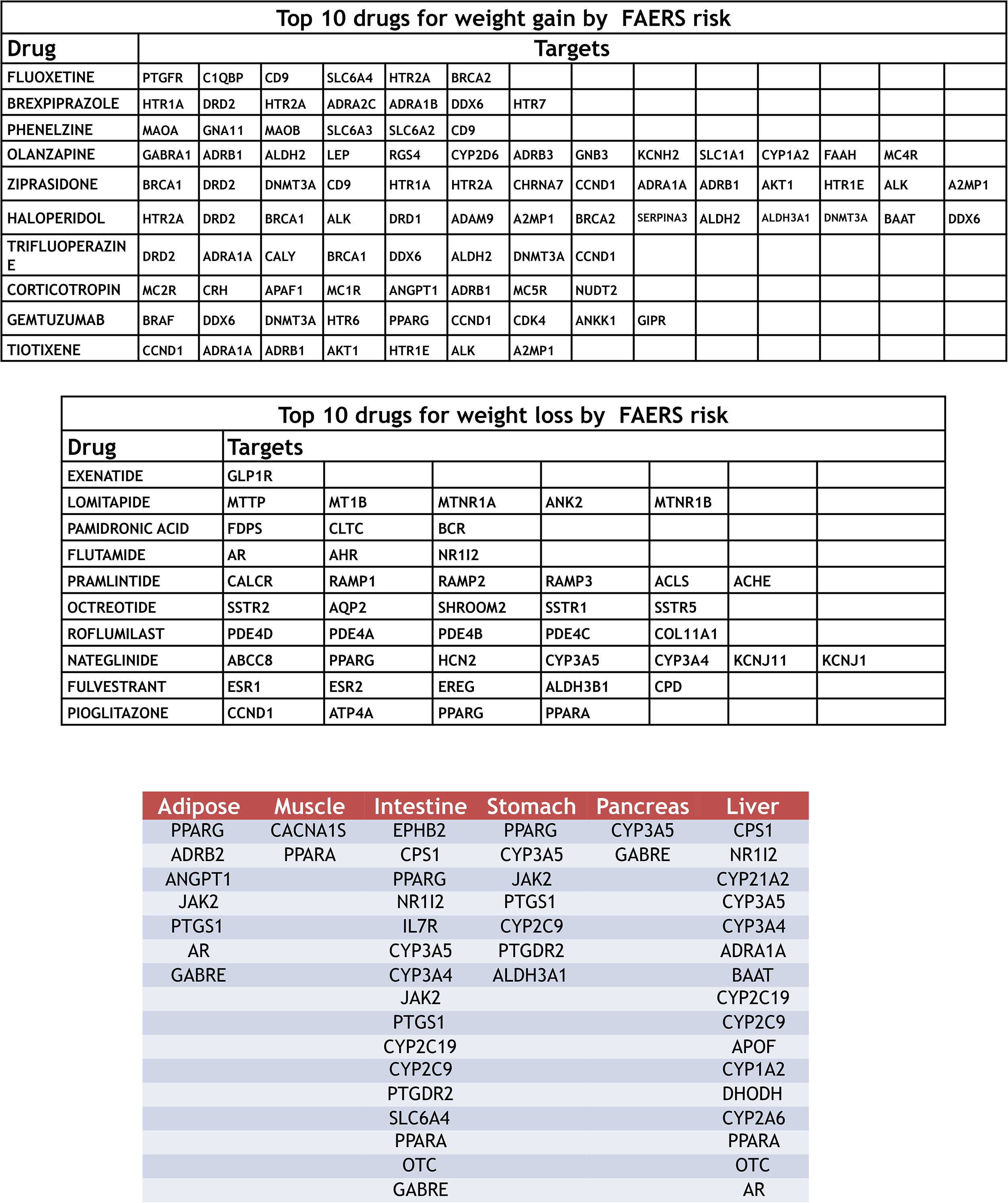
The 10 FDA-approved drugs that have the highest FAERS risk for weight gain and the 10 with the highest risk for weight loss as side effects, and their respective targets. Targets are listed by protein name and were obtained from multiple databases: DrugBank, The Druggable Genome, and Therapeutic Targets Database. See **Supplementary File 5** for the complete list of all the drugs and their respective targets that have weight gain or weight loss as side effects **Organ/tissue distribution of protein targets of weight gain- & weight loss-inducing drugs.** Those targets of the 20 FDA-approved drugs listed in table 1 which are expressed in the 6 peripheral organs/tissues of interest are listed according to their expression retrieved from the GTEx database.

**Figure 7.**
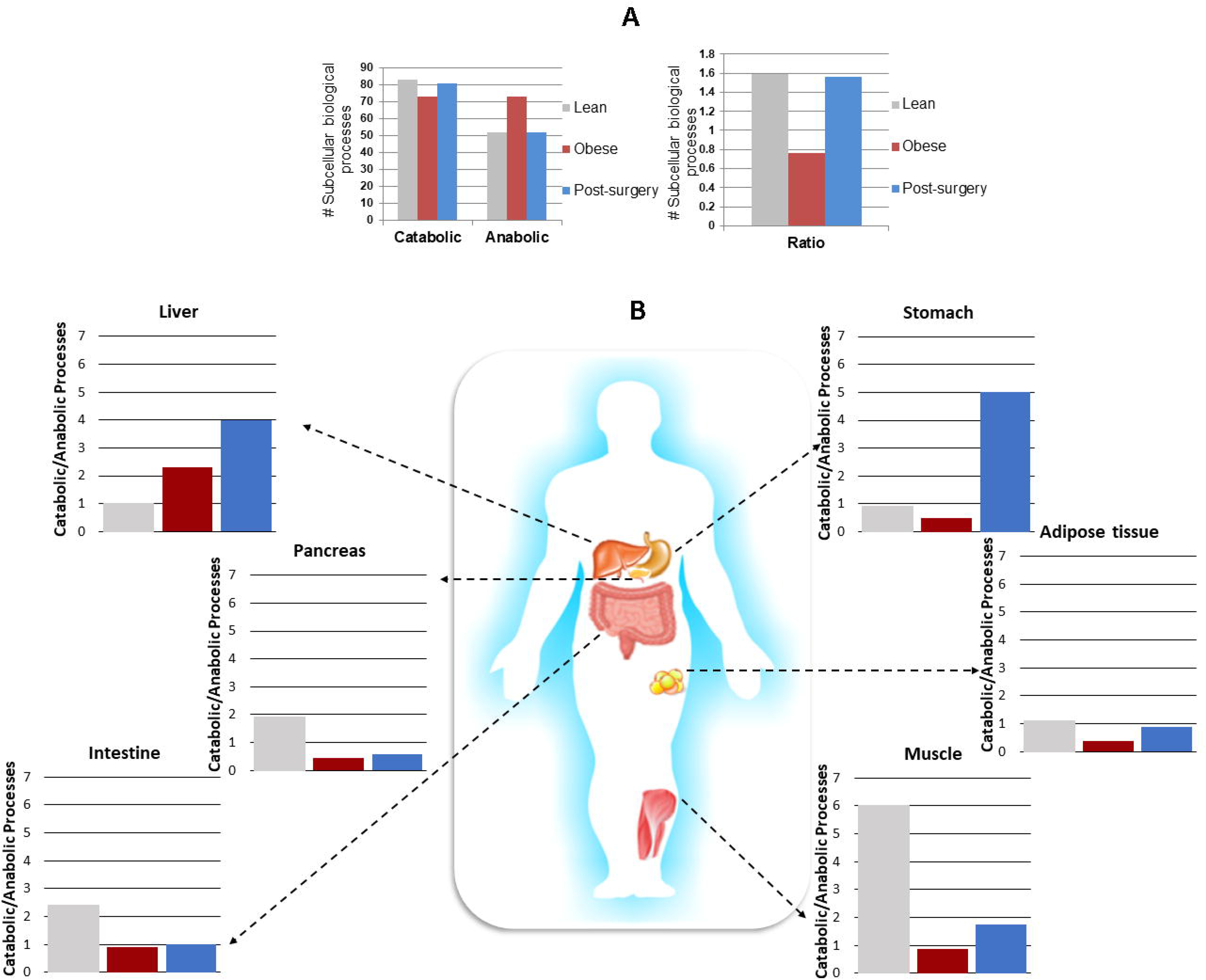
An initial human obesity atlas reveals organ- and tissue-specific metabolic differences in lean, obese, and post-bariatric surgery conditions. **A) Statistically significant catabolic and anabolic processes in lean, obese, and post-surgery subjects.** (Top) The number of catabolic and anabolic processes within the top 100 biological subcellular processes was identified by the MCBO algorithm for the whole body for each physiological condition. (B**ottom**) Ratio of catabolic to anabolic processes for lean, obese and post-bariatric surgery patients. **B) Mapping the ratio of catabolic to anabolic processes onto individual tissues/organs in lean, obese, and post-bariatric surgery subjects.** Using the obesity atlas, differences in overall metabolism can be mapped to individual tissues and organs in persons with different physiological phenotypes.

## Discussion

This atlas, to the best of our knowledge, is among the first to contain most obesity-related genetic, genomic, transcriptomic and physiological data and to show their interactions focusing on non-CNS targets. Our aim in developing the atlas is to obtain new knowledge on the pathophysiology of obesity and to help catalyze new drug-based therapeutic approaches for treating obesity. Most published atlases have focused on molecular data such as gene expression profiles [38, 39] or relationships between gene expression and pathology [40, 41]. Typically, the physiology of obesity has been approached by analyzing one or a few hormones levels in the blood at a time [8, 21-–29], not adequately taking into consideration the complexity of the disease. A key finding from our atlas is that multiple tissues/organs are likely to have significant involvement in obesity, and it may not be possible to rank them and the potential drug targets within them in a simple manner.

Our bottom-up integrative analysis offers multiple insights of the biological organization combining genetic and genomic determinants with different types of physiological dynamics. Several features of our method illustrate how it can be transferable to the study of other systemic diseases. First, it utilizes publicly available knowledge and allows scaling the information by systematic statistical analysis. Second, it allows the integration of datasets: static and dynamic, unbiased, and specified. Third, it permits prediction for tissue/organ-selective new combinatorial drug targets that can be further experimentally tested. We took data from several biological levels, from molecular whole genomic data to cell-level transcriptomic data, to organ-level endocrine data, in order to exhaustively characterize 3 different phenotypic states: lean, obese, and post-bariatric surgery. The integration achieved by connecting dynamical models to network models enables us to identify subcellular processes within the organs of interest. It should be noted that PPI networks allow for the identification of intermediate genes or gene products that could be potentially overlooked [42]. The computational model of the hormone levels and the regression analysis allow for the study of a system from a dynamic perspective and for ranking cell-level information. The use of transcriptomic expression profiles for various organs (**Figure 5**) provides a facile approach to parse the physiological state-selective molecular networks into tissue/organ-based physiological state-selective networks. Enrichment analysis of subcellular biological processes permits us to identify the interactions between the different datasets from a functional perspective. This ability to resolve the molecular networks provides insight into the different cellular pathways related to obesity that are operational in different tissues and organs. While the role of the central nervous system as a target for control of body weight [43] has been well studied in terms of hunger circuits [44], similar information regarding other tissues and organs have not been available until now. Our analysis unravels the complexity underlying obesity using a systems biology approach: genomic determinants widely distributed on all the non-sex chromosomes (**Figure 2**) and cellular pathways that are operational in many tissues and organs (**Figure 7**, **Supplementary Figure S14**). From the perspective of physiological states, our findings provide a clear representation of the divergence of the plasma level of several metabolic hormones and of the catabolic and anabolic subcellular processes among lean, obese and post-bariatric surgery patients in six tissues and organs. In this context, skeletal muscle, pancreas, intestine and adipose tissue emerged as the most perturbed (compared to the lean state) by the obese state. Stomach, liver, pancreas and skeletal muscle seem to be the most perturbed (compared to the lean state) in the post-bariatric surgery state. Thus, it appears that the post-bariatric state is different from the lean state in terms of organ function, although they are phenotypically similar. We believe this represents the first identification of the perturbation of pathways implicated in energy production and expenditure in obesity by data integration. Additionally, from the atlas, we can deduce by unbiased analysis that in lean people catabolic processes predominate while in obese people anabolic processes predominate. Although this conclusion is intuitively obvious, have it established by unbiased information and analysis of multiple datasets, our findings provide a clear representation of the divergence of the level of several energy-regulating hormones and of the divergence in the catabolic and anabolic subcellular processes among lean, obese and post-bariatric surgery patients. According to our analysis, these alterations stem primarily from perturbed transport and mobilization of cholesterol and LDL, glycolysis, and glycogen synthesis (**Supplementary Figure S14**). Despite this relatively simple overall conclusion, the distribution of these anabolic and catabolic processes among multiple organs highlights the underlying complexity in going from physiological knowledge to drug development. Adipose tissue, intestine and, rather unexpectedly, stomach seem to be the most promising organs for new potential drug targets (Table 2), although we cannot fully discount whether the use of data from bariatric surgery influences our ranking of the stomach. According to our results, PPARG (peroxisome proliferator-activated receptor gamma) seems to be significant in adipose tissue and intestine, and PPARA (peroxisome proliferator-activated receptor alpha) in muscle, intestine, and liver. This is consistent with their roles in glucose metabolism and fatty acid storage. ADBR2 (the beta-2-adrenergic receptor) which is known to participate in glycogenesis and gluconeogenesis in the liver and skeletal muscle, seems to play a role in adipose tissue as well. In the intestine, the presence of IL7R (interleukin7 receptor) and PTGDR2 (prostaglandin DP2 receptor) could be an indication of inflammatory states, which have been shown to be correlated with obesity [3]. ALDH3A1 (aldehyde dehydrogenase 3 family member A1) is involved in lipid peroxidation and could represent an interesting target for further investigation. While the presence of several genes of the cytochrome family is to be expected in the liver, our results suggest that APOF (apolipoprotein F) involved in the transport of cholesterol, could be considered for future therapeutic approaches.

**Table 1:**
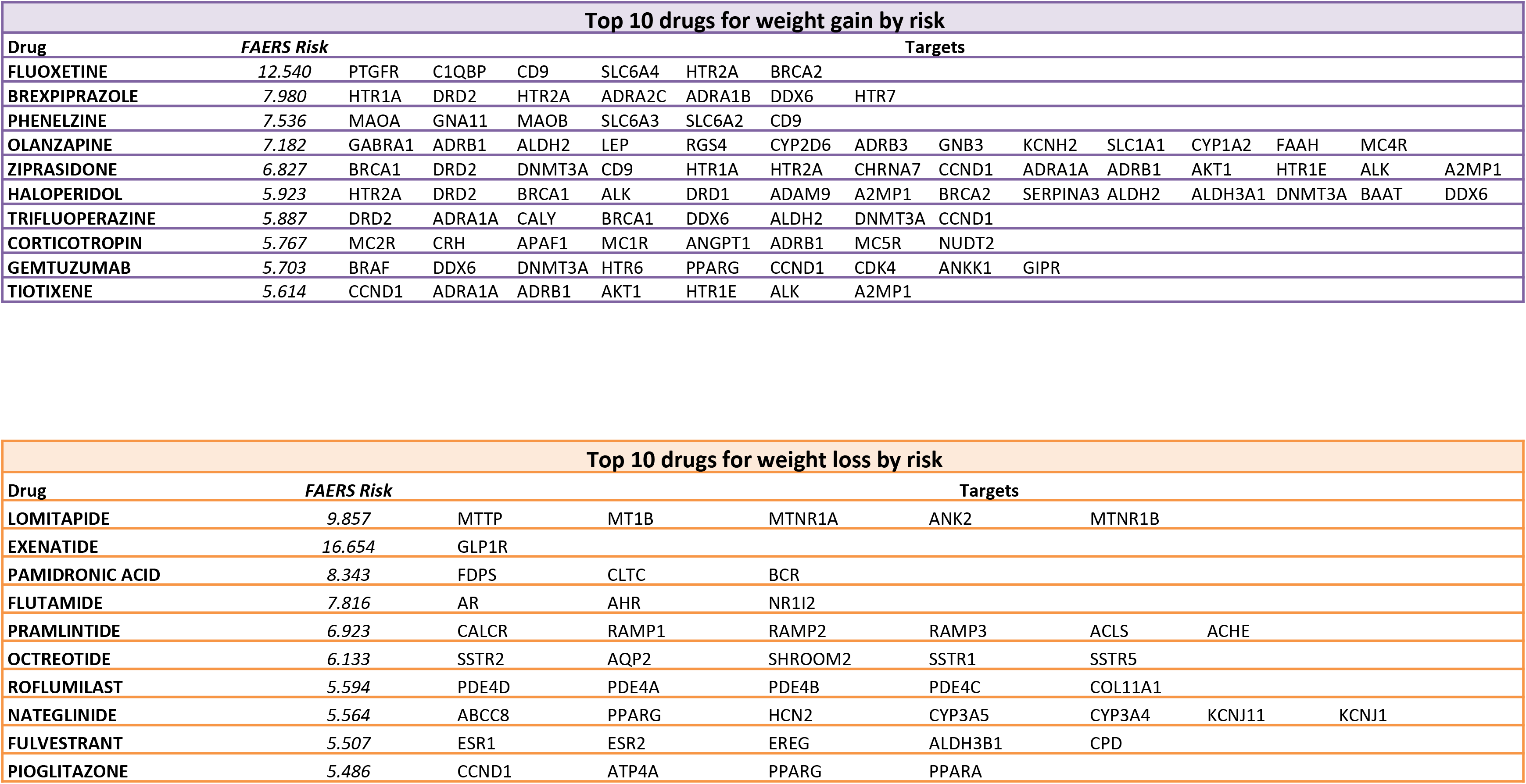

**Table 2:**

| Adipose | Muscle | Intestine | Stomach | Pancreas | Liver |
| --- | --- | --- | --- | --- | --- |
| PPARG | CACNA1S | EPHB2 | PPARG | CYP3A5 | CPS1 |
| ADRB2 | PPARA | CPS1 | CYP3A5 | GABRE | NR1I2 |
| ANGPT1 |  | PPARG | JAK2 |  | CYP21A2 |
| JAK2 |  | NR1I2 | PTGS1 |  | CYP3A5 |
| PTGS1 |  | IL7R | CYP2C9 |  | CYP3A4 |
| AR |  | CYP3A5 | PTGDR2 |  | ADRA1A |
| GABRE |  | CYP3A4 | ALDH3A1 |  | BAAT |
|  |  | JAK2 |  |  | CYP2C19 |
|  |  | PTGS1 |  |  | CYP2C9 |
|  |  | CYP2C19 |  |  | APOF |
|  |  | CYP2C9 |  |  | CYP1A2 |
|  |  | PTGDR2 |  |  | DHODH |
|  |  | SLC6A4 |  |  | CYP2A6 |
|  |  | PPARA |  |  | PPARA |
|  |  | OTC |  |  | OTC |
|  |  | GABRE |  |  | AR |

Overall, our analysis indicates that drug treatment of obesity aimed at non-CNS targets is most likely to be successful if it is pleiotropic in nature, affecting cellular regulation at multiple sites.

## Online Methods

### Human genetic dataset

The human genetic dataset is composed of genes selected from the NCBI Gene Database by text mining. Using “*trait* + [orgn] homo sapiens” as a query term we retrieved genes that were associated with 12 terms related to obesity according to published literature as of May 2018: “obesity”, “obese”, “bariatric surgery”, “Body Mass Index OR BMI”, “fat”, “fat mass”, “fat free mass OR FFM”, “lean OR leanness”, “overweight”. A cut-off of at least 2 publications per gene was set. We retrieved 2734 overall unique genes. The trait-specific and complete list of genes can be consulted in the **Supplementary File 1**.

### Human genomic dataset

To create this dataset we collected genomic data available in the Genome Wide Association Studies Catalog [3, 45] from the May 2017 version. The GWAS data are currently mapped to Genome Assembly GRCh38.p7 and dbSNP Build 147. From the list of selectable diseases and traits in the catalog we selected 11 terms of interest for our study with their respective list of SNPs and associated genes: adiposity, body fat percentage, body mass (lean), body mass index, fat body mass, obesity, visceral adipose tissue/subcutaneous adipose tissue ratio, waist circumference, waist-hip ratio, weight, weight loss (gastric bypass surgery). The respective numbers of SNPs, intergenic and intragenic regions are listed in **Supplementary Table S3**. The results were filtered to exclude studies that didn’t provide replication; only adult subject cohort studies were considered; finally, only SNPs that were associated with traits with p-value ≤ *5*10*^-8^, the standard genomic statistical significance threshold, were selected. A total of 751 SNP-trait associations, 220 mapped genes, and 30 intergenic regions were found. To obtain a list of unique SNPs, if a SNP was reported in more than one study, only the most significant SNP-trait association score (smallest p-value) was considered. The GWAS catalog already provides genome mapping for the listed genes. Nevertheless, the mapping was further manually curated utilizing Genome Browser [15] to assess any new potential mapping though ENCODE [14]. The final curated list of SNP-trait associations for each trait of interest is stored in **Supplementary File 2**.

### Protein-Protein interaction network analysis

We utilized X2K software [46] to create a protein-protein interaction (PPI) network. X2K provides a large-scale protein-protein interaction network made from 18 sources that connect seed lists of genes/proteins. The user can map a list of seed genes to a human interactome. We utilized the 2631 genes in the genetic and genomic datasets as seeds to build a PPI network. A path length of 2 was chosen meaning that the path between two seed nodes requires two edges to connect the nodes, so there is only one intermediate protein between them, i.e., a protein that interacts with both seed nodes. BIND [47], HPRD [48], MINT [49], Biocarta [50], InnateDB [51], MIPS [52], SNAVI [53], BioGRID [54], INTact [55], DIP [56], PDZbase [57], and KEGG [58] databases were selected to extrapolate the intermediate genes from the human interactome. 4637 intermediates were identified.

### Multicompartmental dynamical model

We built multicompartmental dynamical models using coupled ordinary differential equations to describe the endocrinological response to glucose ingestion in a lean, obese, and post-bariatric surgery human subject by modeling the plasma levels of eleven hormones associated with the regulation of satiation and appetite. We built a model for each hormone and used the same glucose stimulus for each. Our model differs from the previously published ones [59, 60] because (a) it describes endogenous instead of exogenous plasma levels of the hormones; (b) it provides the kinetic behavior of eleven hormones; (c) it takes into account multiple compartments and organs; (d) it describes three different physiological states; (e) the aim of the model is to identify a set of kinetic parameters that best describe the temporal progression of the experimentally observed values. These parameters would then allow us to determine statistically significant differences between lean, obese and post-bariatric surgery endocrinology. These parameters can then be varied in an unbiased manner to identify the fragile and robust parameters of the system as whole.

### Collation of the experimental data of hormone plasma levels

Data points for each individual time course were obtained from published data [8, 22–32]. Inclusion criteria for the published data were: a) The selected hormone must be associated to the regulation of appetite level and satiation level in human subjects; b) The plasma level of the hormones must be endogenous; c) Time points must cover at least one hour of a fasting state and two hours of a postprandial state; d) Subjects should not be affected by any comorbidity; d) Postsurgery subjects must have undergone Roux Y Gastric Bypass and should preferably be one year post-surgery.

Data points for lean, obese and post-RYGB surgery were collected for 3 orexigenic hormones (ghrelin, adiponectin, and glucagon) and eight anorexigenic hormones (leptin, insulin, amylin, cholecystokinin, peptide YY, glucagon-like peptide 1, gastric inhibitory peptide-1, oxyntomodulin) (**Supplementary Table 4**). Each point in every time course is the mean ± S.E.M. of time points from different published data (A complete list of the time points is provided in **Supplementary File 3**). All the different units of concentration of hormones were converted to pmol/L utilizing the molecular weight of each hormone.

### Compartmental dynamic ordinary differential equation (ODE) model

Our model describes the kinetics of eleven different hormones secreted by the different organs in the digestive system in response to meal ingestion. Glucose digestion, absorption, production, utilization and degradation and each hormone’s secretion, first pass through the liver, utilization and degradation were considered. The model comprises seven explicit compartments (stomach, intestine, pancreas, adipocytes, plasma, muscle and liver) and one black-box compartment (degradation). A scheme of the interactions in the model is shown in **Figure 3A**. Given the complexity of the system, and the availability of only plasma glucose and hormone concentrations to find the parameters, the parameter estimation methodologies were used to build a reliable model [61–64]. The model parameters were numerically identified by nonlinear least-squares as implemented in MATLAB 2018a.

### Glucose Kinetics

#### Endogenous Glucose Production (EGP)

The functional description of EGP in terms of glucose and insulin signals comprises a direct glucose signal and both delayed and anticipated insulin signals

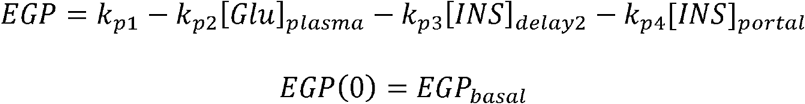

where [*Glu*]_*plasma*_ is the concentration of the plasma glucose, [*INS*]_*portal*_ is the concentration of insulin in the portal vein, [*INS*]_*delay*2_ [*INS*]_delay_ is a delayed insulin signal realized with a chain of two compartments:

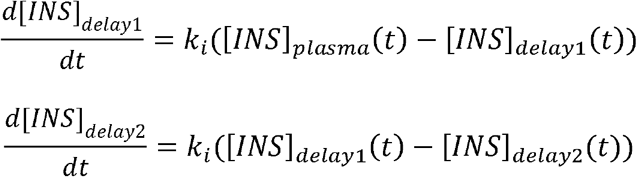

*k*_*p*1_ (mg/kg/min) is the basal rate of EGP at zero glucose and insulin, *k*_*p*2_ (min^-1^) is the parameter for liver glucose effectiveness in reducing plasma glucose, *k*_*p*3_ (mg/kg/min/(mol/l)) is the parameter governing amplitude of insulin action on the liver, *k*_*p*4_ (mg/kg/min/(mol/kg)) is the parameter for amplitude of portal insulin action on the liver and *k_i_* is the parameter accounting for delay between the insulin signal and insulin action. EGP is also constrained to be nonnegative.

#### Glucose Ingestion and Absorption

Glucose transits through the stomach and intestine by assuming the stomach to be represented by two compartments (one for solid [*Glu*]_*stomach1*_, and one for triturated phase [*Glu*]_*stomach*2_), while a single compartment is used to describe the intestine. Glucose concentration in the stomach as a function of time is given by:

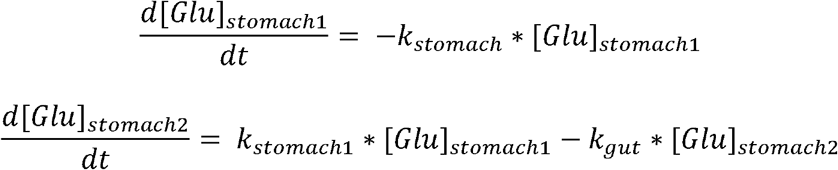

Glucose concentration in the intestine is given by:

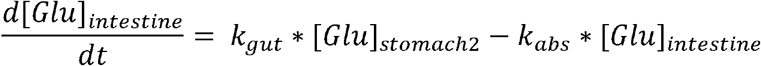

#### Glucose Utilization

Hormone-independent utilization represents glucose uptake by the brain and erythrocytes *F_cns_*(*t*). Peripheral tissue hormone-dependent utilization depends non-linearly on glucose in the tissues and it is described by a Michaelis–Menten kinetics:

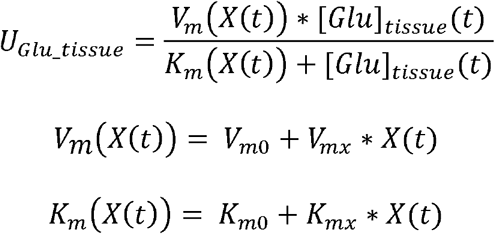

As we have 11 hormones:

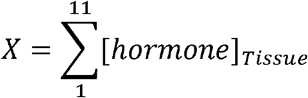

Total glucose utilization *U*_*Glu*_:

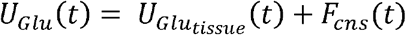

### Hormone Kinetics

Glucose ingestion triggers the secretion of orexigenic and anorexigenic hormones from different peripheral organs and tissue.

#### Hormone Secretion

Hormone secretion into plasma by specific organs occurs if glucose concentration in that organ exceeds a certain threshold. Hormone-specific secretions are described below.

#### Stomach Secretion of Ghrelin (GHRL)

If 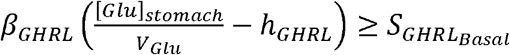 then ghrelin secretion is triggered:

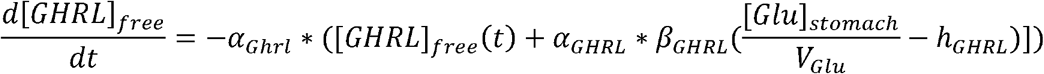

otherwise

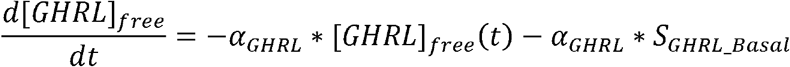

Where *α_GHRL_* (min^-1^) is the delay between the glucose signal and ghrelin secretion, *β_GHRL_* (pmol/kg/min per mg/dL is the stomach “X/A cells” responsivity to glucose, and *h_GHRL_* (mg/dL) is the threshold level of glucose above which the “X/A cells” initiate to produce ghrelin (has been set to the basal glucose concentration to guarantee system steady state in basal condition).

#### Intestine Secretion of Glucagon-like Peptide 1 (GLP-1)

If 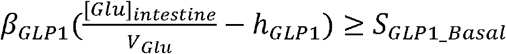 then GLP-1 secretion is triggered:

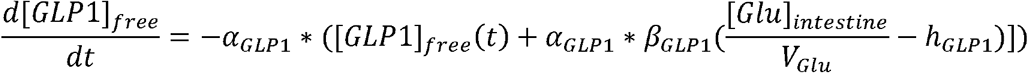

otherwise

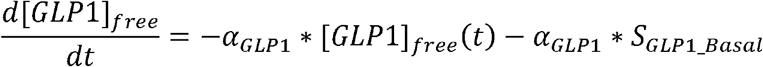

Where *α*_*GLP*1_ (min^-1^) is the delay between glucose signal and GLP-1 secretion, *h*_*GLP*1_ (pmol/kg/min per mg/dL) is the lower intestine L cells responsivity to glucose, and *h*_*GLP*1_ (mg/dL) is the threshold level of glucose above which the L cells begin to produce GLP1 (has been set to the basal glucose concentration to guarantee system steady state in basal condition).

#### Intestine Secretion of Peptide YY (PYY)

If 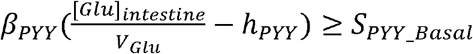 then PYY secretion is triggered:

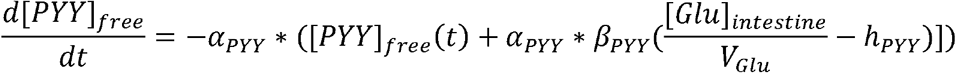

otherwise

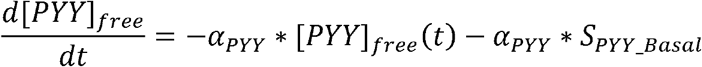

Where *α_PYY_* (min^-1^) is the delay between glucose signal and PYY secretion, *β_PYY_* (pmol/kg/min per mg/dL) is the lower intestine L cells responsivity to glucose, and *h_PYY_* (mg/dL) is the threshold level of glucose above which the L cells begin to produce PYY (has been set to the basal glucose concentration to guarantee system steady state in basal condition).

#### Intestine Secretion of Gastric Inhibitor Peptide (GIP)

If 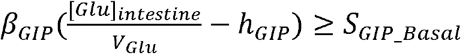 then GIP secretion is triggered:

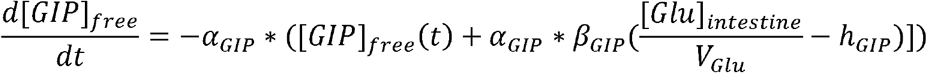

otherwise

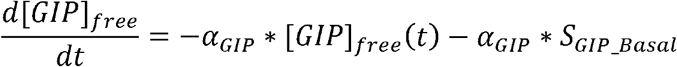

Where *α_GIP_* (min^-1^) is the delay between glucose signal and GIP secretion, *β_GIP_* (pmol/kg/min per mg/dL) is the lower intestine L cells responsivity to glucose, and *h_GIP_* (mg/dL) is the threshold level of glucose above which the L cells begin to produce GIP (has been set to the basal glucose concentration to guarantee system steady state in basal condition).

#### Intestine Secretion of Oxyntomodulin (OXM)

If 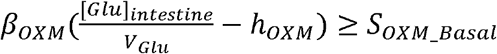 then cholecystokinin secretion is triggered:

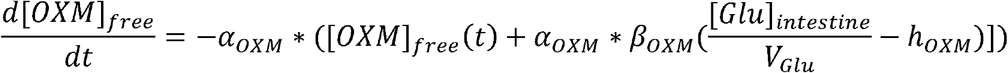

otherwise

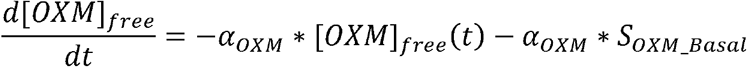

Where *α_OXM_* (min^-1^) is the delay between glucose signal and OXM secretion, *β_OXM_* (pmol/kg/min per mg/dL) is the lower intestine L cells responsivity to glucose, and *h_OXM_* (mg/dL) is the threshold level of glucose above which the L cells begin to produce OXM (has been set to the basal glucose concentration to guarantee system steady state in basal condition).

#### Intestine Secretion of Cholecystokinin (CKK)

If 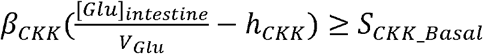 then cholecystokinin secretion is triggered:

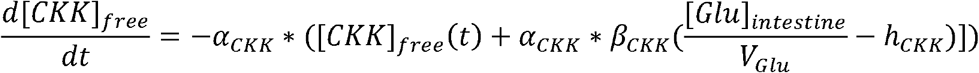

otherwise

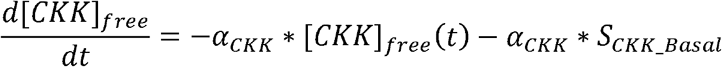

Where *α_CKK_* (min^-1^) is the delay between glucose signal and CKK secretion, *β_CKK_* (pmol/kg/min per mg/dL) is the lower intestine I cells responsivity to glucose, and *h_CKK_* (mg/dL) is the threshold level of glucose above which the I cells begin to produce CKK (has been set to the basal glucose concentration to guarantee system steady state in basal condition).

#### Pancreatic secretion of insulin (INS)

If 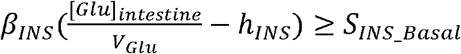 then insulin secretion is triggered:

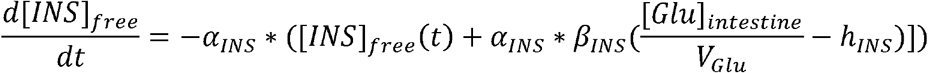

otherwise

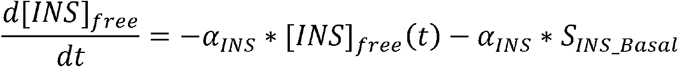

Where *α_INS_* (min^-1^) is the delay between glucose signal and insulin secretion, *β_INS_* (pmol/kg/min per mg/dL) is the pancreatic β cells responsivity to glucose, and *h_INS_* (mg/dL) is the threshold level of glucose above which the β cells begin to produce insulin (has been set to the basal glucose concentration to guarantee system steady state in basal condition).

#### Pancreatic Secretion of Amylin (AMY)

If 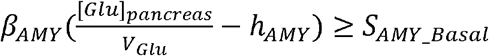 then amylin secretion is triggered:

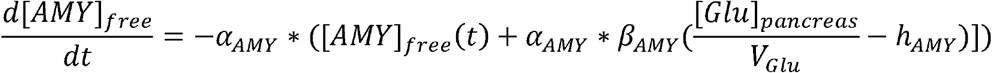

otherwise

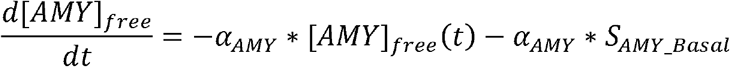

Where *α_AMY_* (min^-1^) is the delay between glucose signal and amylin secretion, β_AMY_ (pmol/kg/min per mg/dL) is the pancreatic β cells responsivity to glucose, and *h_AMY_* (mg/dL) is the threshold level of glucose above which the β cells begin to produce amylin (has been set to the basal glucose concentration to guarantee system steady state in basal condition).

#### Pancreatic Secretion of Glucagon (GCG)

If

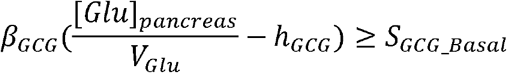

#### Glucagon secretion is triggered

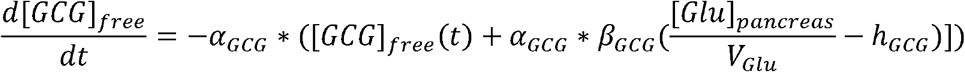

otherwise

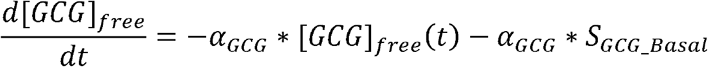

Where *α_GCG_* (min^-1^) is the delay between glucose signal and glucagon secretion, β_GCG_ (pmol/kg/min per mg/dL) is the pancreatic α cells responsivity to glucose, and *h_GCG_* (mg/dL) is the threshold level of glucose above which the α cells begin to produce glucagon (has been set to the basal glucose concentration to guarantee system steady state in basal condition).

#### Adipose Tissue Secretion of Leptin (LEP)

If 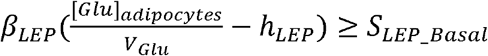 then leptin secretion is triggered:

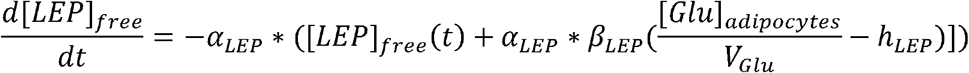

otherwise

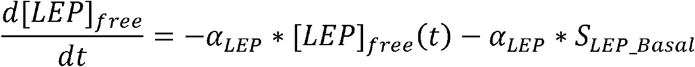

Where *α_LEP_* (min^-1^) is the delay between glucose signal and leptin secretion, *β_LEP_* (pmol/kg/min per mg/dL) is the adipocytes responsivity to glucose, and *h_LEP_* (mg/dL) is the threshold level of glucose above which the adipocytes begin to produce leptin (has been set to the basal glucose concentration to guarantee system steady state in basal condition).

#### Adipose Tissue Secretion of Adiponectin (ADIPOQ)

If 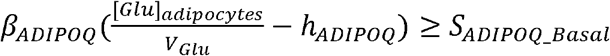 then adiponectin secretion is triggered:

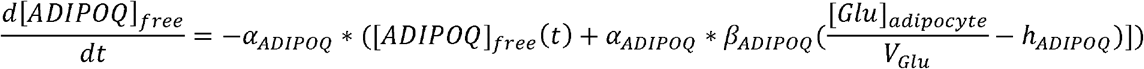

otherwise

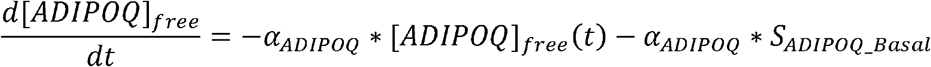

Where *α_ADIPOQ_* (min^-1^) is the delay between glucose signal and adiponectin secretion, *β_ADIPOQ_* (pmol/kg/min per mg/dL) is the adipocytes responsivity to glucose, and *h_ADIPOQ_* (mg/dL) is the threshold level of glucose above which the adipocytes initiate to produce adiponectin (has been set to the basal glucose concentration to guarantee system steady state in basal condition).

#### Degradation and Plasma Level of the Hormones

Degradation occurs both in the liver (first pass) and in the periphery and it follows the kinetics listed below.

#### Ghrelin Degradation and Plasma Level

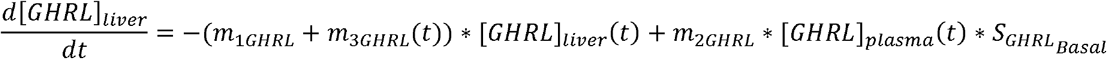

Which determines the temporal changes of the plasma level of GHRL:

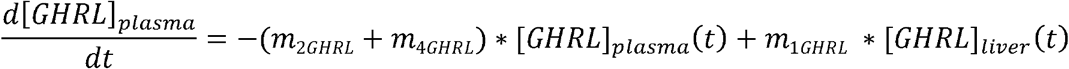

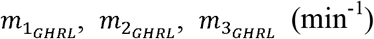 are rate parameters. 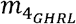 is peripheral degradation. Hepatic extraction (HE) that determines liver degradation of ghrelin is described by:

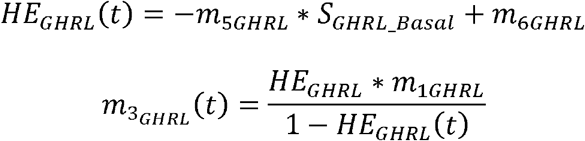

At basal steady state:

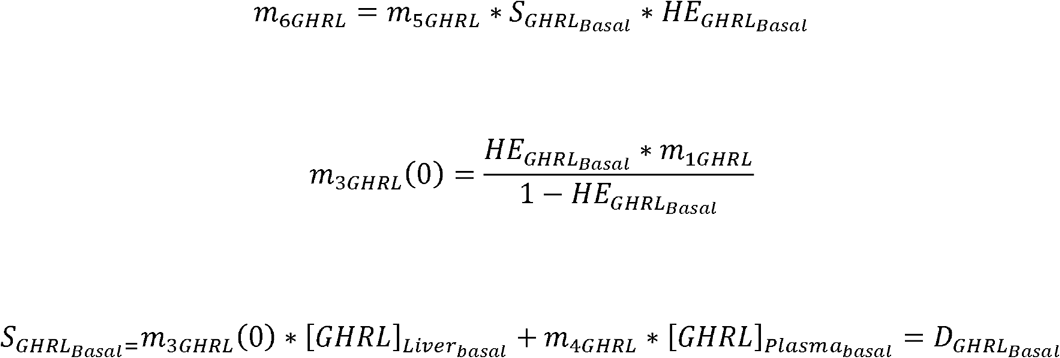

where *D_GHRL_Basa1__* is the basal total degradation.

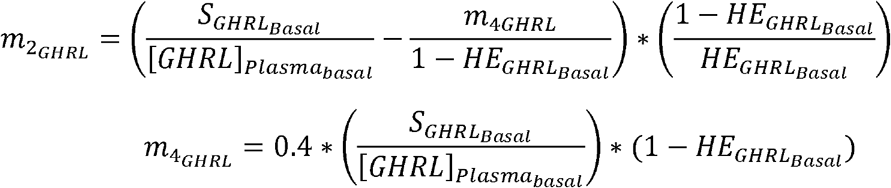

#### GLP-1 Degradation and Plasma Level

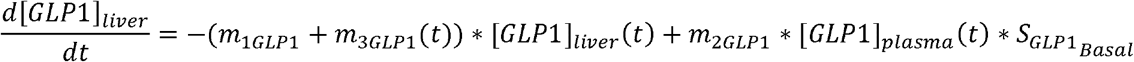

Which determines the temporal change in the plasma level of GLP1:

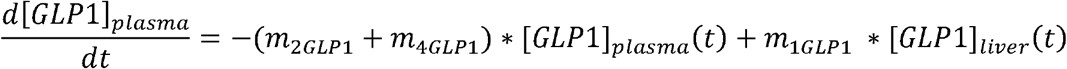

*m*_1*GLP*1_, *m*_2*GLP*1_, *m*_3+*GLP*1_ (min^-1^) are rate parameters. *m*_4*GLP*1_ is peripheral degradation. Hepatic extraction (HE) that determines liver degradation of ghrelin is described by:

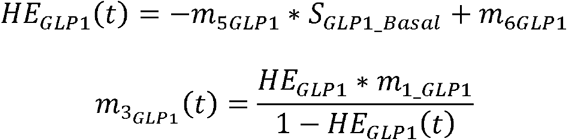

At basal steady state:

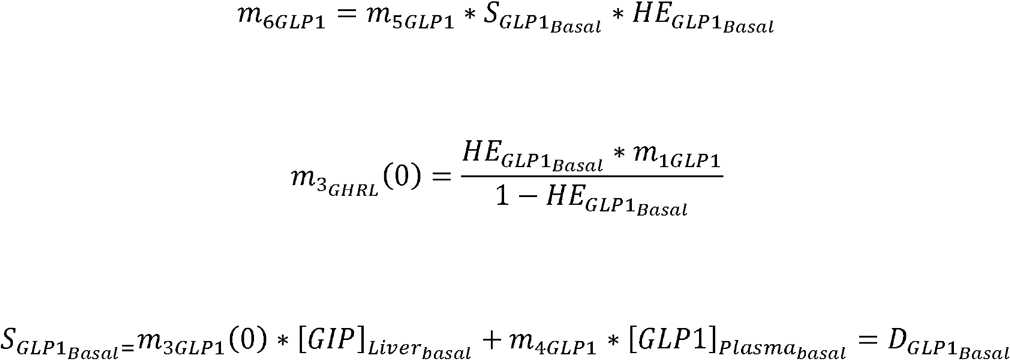

where D_GLP1basa1_ is the basal total degradation.

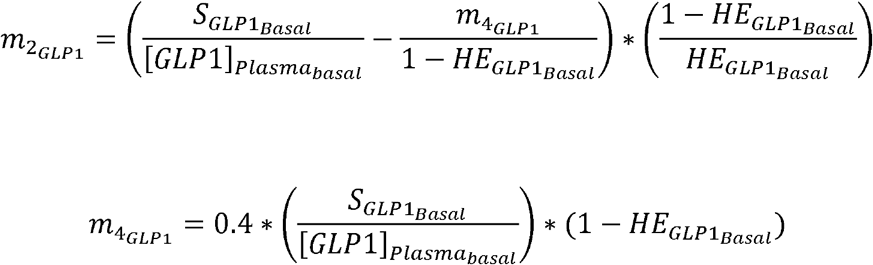

#### PYY Degradation and Plasma Level

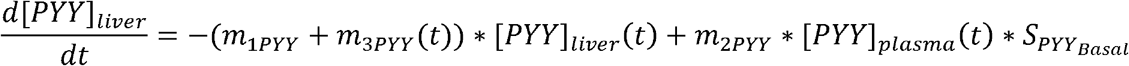

Which determines the temporal changes in the plasma level of PYY:

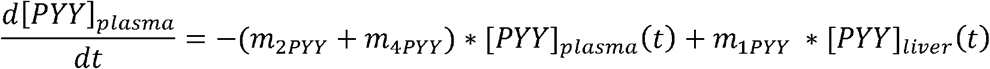

*m*_1*PYY*_, *m*_2*PYY*_ and *m*_4*PYY*_ (min^-1^) are rate parameters. *m*_4*PYY*_ is peripheral degradation. Hepatic extraction (HE) that determines liver degradation of PYY is described by:

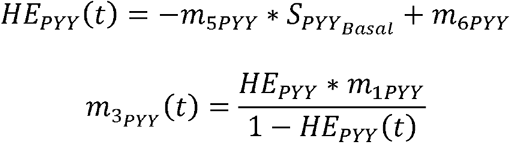

At basal steady state:

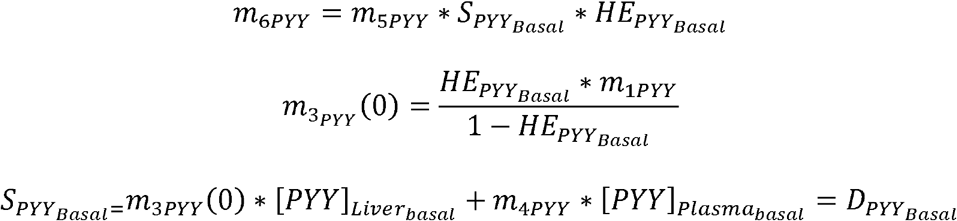

where *D_PYY_Basal__* is the basal total degradation.

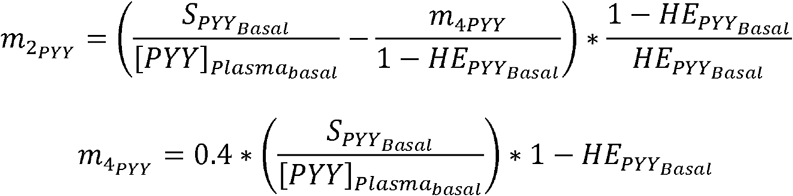

#### GIP Degradation and Plasma Level

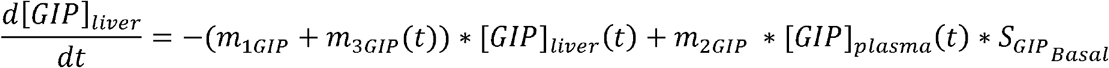

Which determines the temporal changes in the plasma level of GIP:

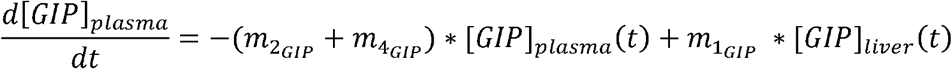

*m*_1*GIP*_, *m*_2*GIP*_, *m*_4*GIP*_ (min^-1^) are rate parameters. *m*_4*GIP*_ is peripheral degradation. Hepatic extraction (HE) that determines liver degradation of GIP is described by:

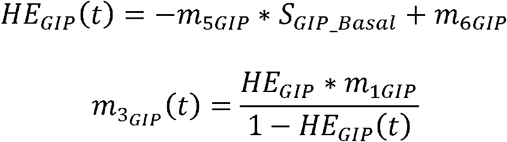

At basal steady state:

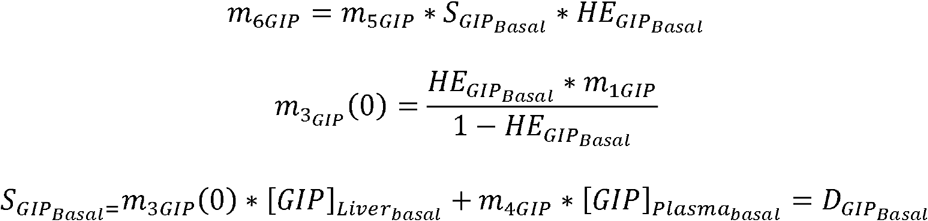

where *D_GIP_Basal__* is the basal total degradation.

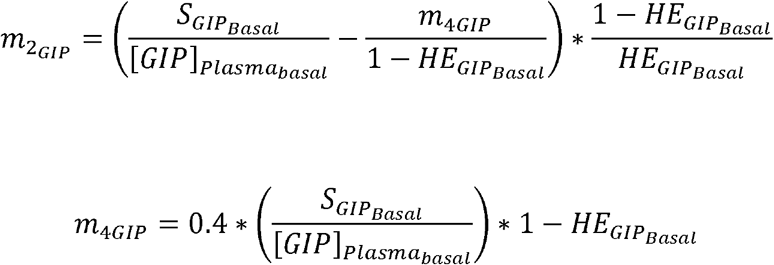

#### OXM Degradation and Plasma Level

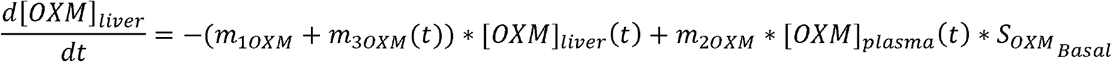

Which determines the temporal changes in the plasma level of OXM:

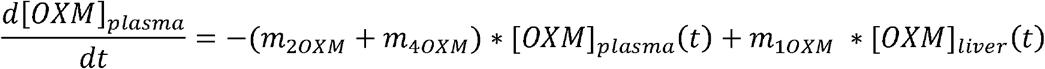

*m*_1*OXM*_, *m*_2*OXM*_, *m*_3*OXM*_ (min^-1^) are rate parameters. *m*_3*OXM*_ is peripheral degradation. Hepatic extraction (HE) that determines liver degradation of OXM is described by:

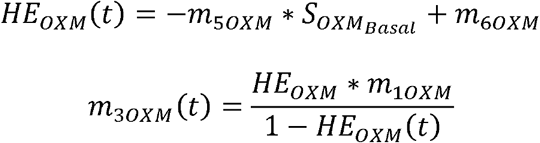

At basal steady state:

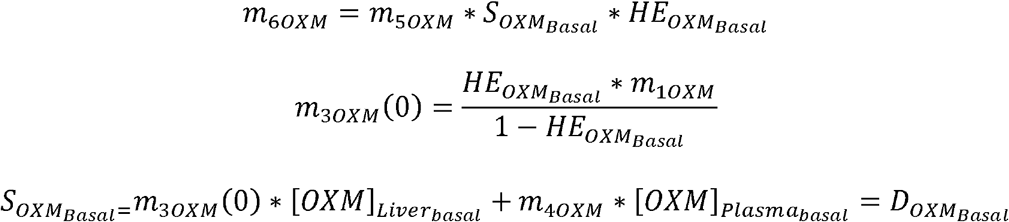

where *D_OXM_Basal__* is the basal total degradation.

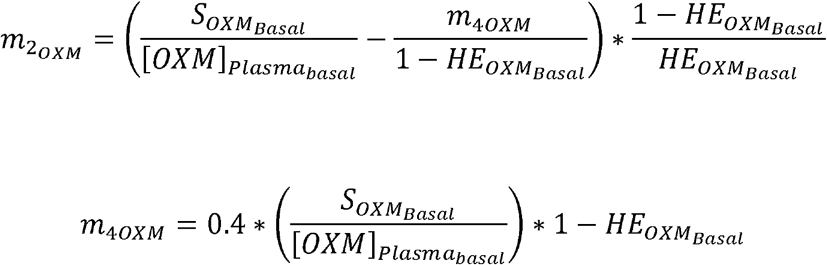

#### CCK Degradation and Plasma Level

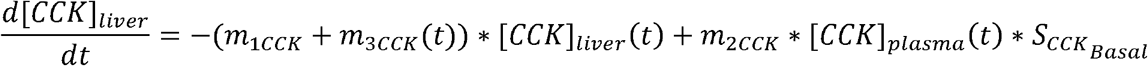

Which determines the temporal changes of the plasma level of CCK:

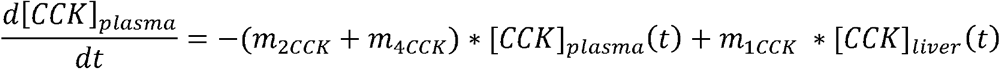

*m*_1*CCK*_, *m*_2*CCK*_, *m*_3*CCK*_ (min^-1^) are rate parameters. *m*_4*CCK*_ is peripheral degradation. Hepatic extraction (HE) that determines liver degradation of CCK is described by:

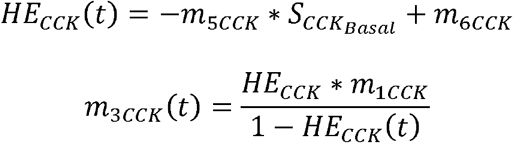

At basal steady state:

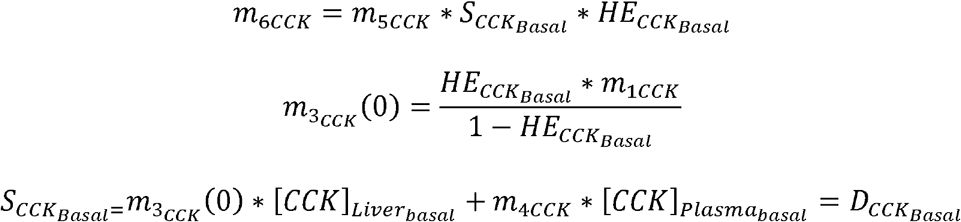

where *D_CCK_Basal__* is the basal total degradation.

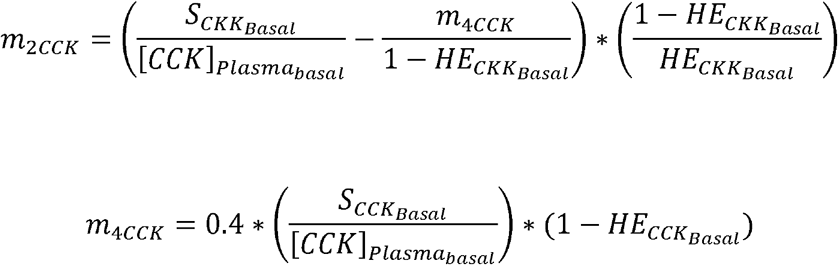

#### Insulin Degradation and Plasma Level

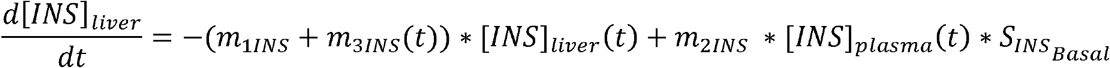

Which determines a plasma level of:

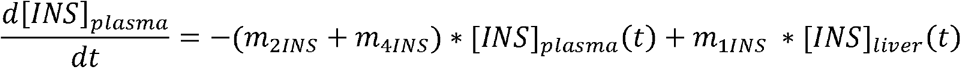

*m*_1*INS*_, *m*_2*INS*_ and *m*_3*INS*_ (min^-1^) are rate parameters. *m*_4*INS*_ is peripheral degradation. Hepatic extraction (HE) that determines liver degradation of insulin is described by:

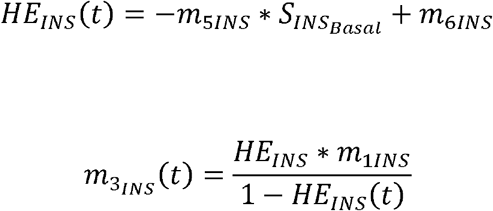

At basal steady state:

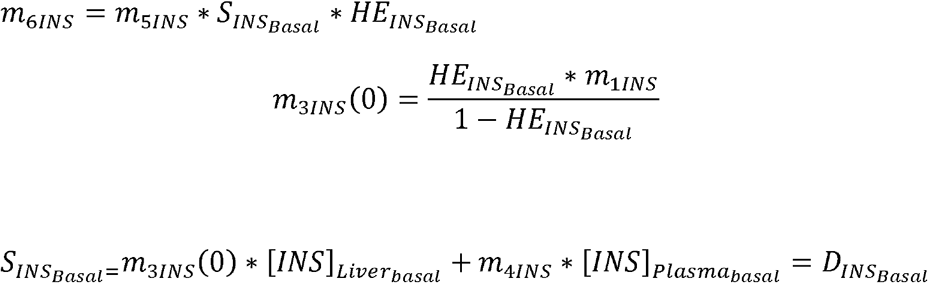

where *D_INS__Basal_* is the basal total degradation.

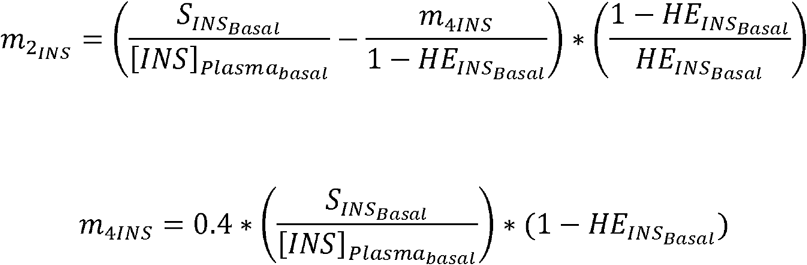

#### Amylin Degradation and Plasma Level

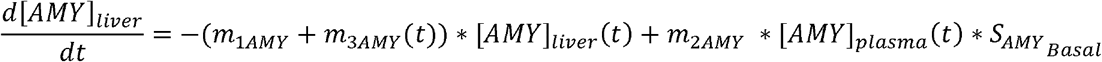

Which determines the temporal changes of the plasma level of amylin:

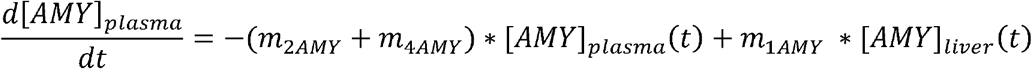

*m*_1*AMY*_, *m*_2*AMY*_ and *m*_3*AMY*_ (min^-1^) are rate parameters. *m*_4*AMY*_ is peripheral degradation. Hepatic extraction (HE) that determines liver degradation of amylin is described by:

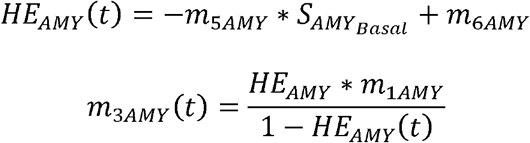

At basal steady state:

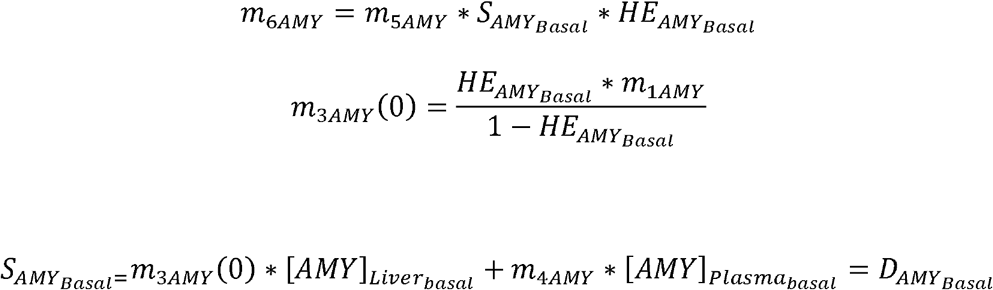

where *D_AMY_Basal__* is the basal total degradation.

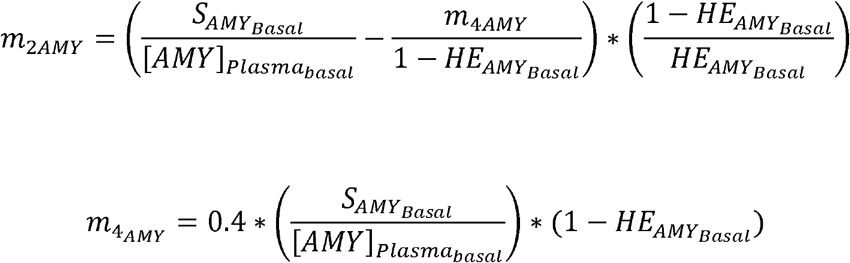

#### Glucagon Degradation and Plasma Level

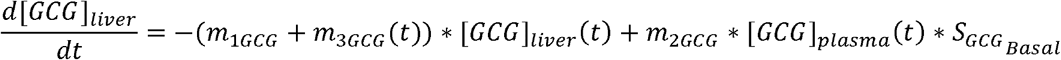

Which determines the temporal changes of the plasma level of glucagon:

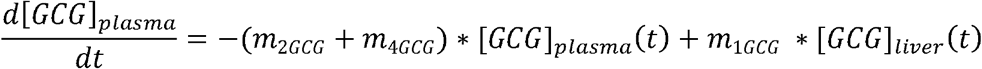

*m*_1*CCG*_, *m*_2*GCG*_ and *m*_3*GGG*_(min^-1^) are rate parameters. *m*_4*GCG*_ is peripheral degradation. Hepatic extraction (HE) that determines liver degradation of glucagon is described by:

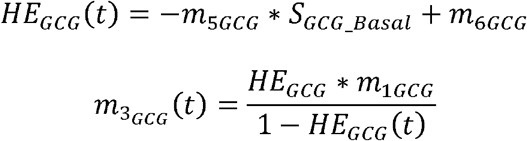

At basal steady state:

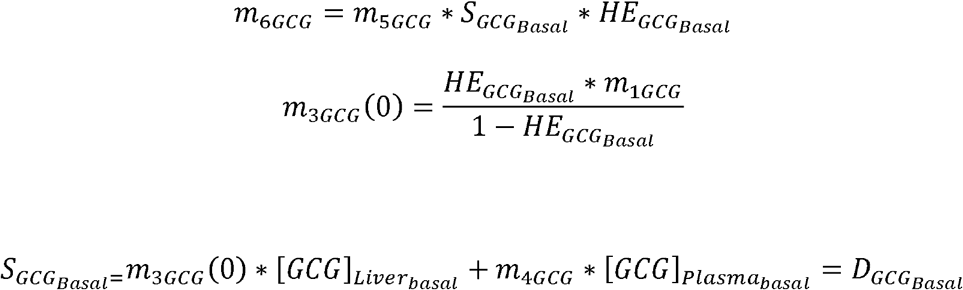

where *D_GCG_Basal__*, is the basal total degradation.

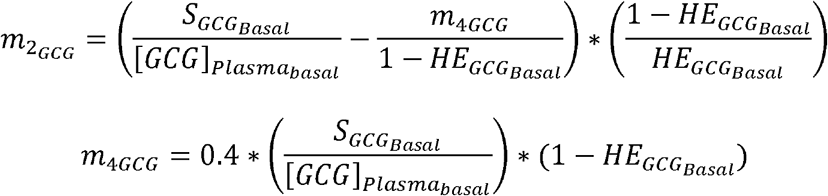

#### Leptin Degradation and Plasma Level

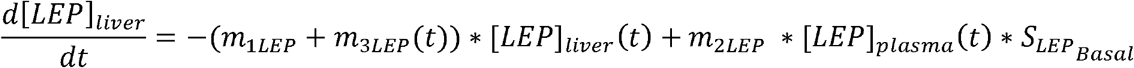

Which determines the temporal changes of the plasma level of leptin:

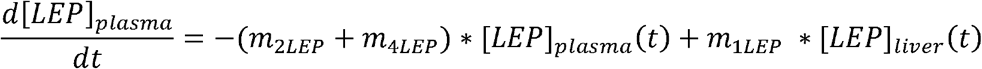

*m*_1*LEP*_, *m*_2*LEP*_ *and m*_3*LEP*_ (min^-1^) are rate parameters. *m*_4*LEP*_ is peripheral degradation. Hepatic extraction (HE) that determines liver degradation of leptin is described by:

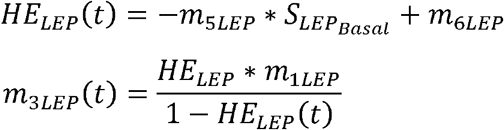

At basal steady state:

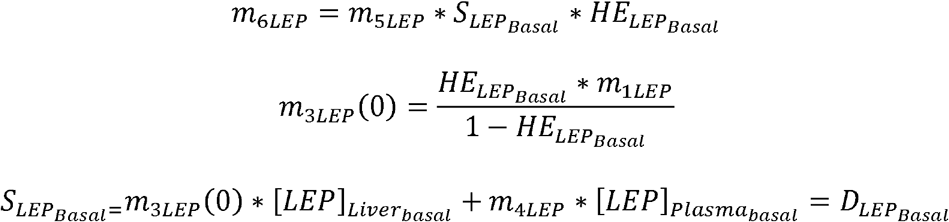

where *D_LEP_Basal__* is the basal total degradation.

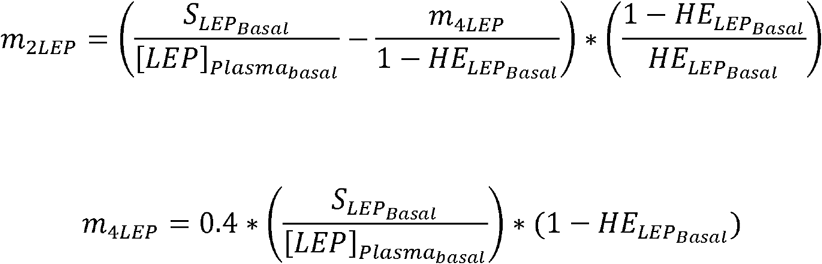

#### Adiponectin Degradation and Plasma Level

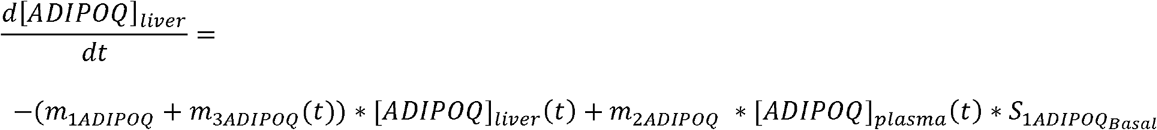

Which determines the temporal changes of the plasma level of adiponectin:

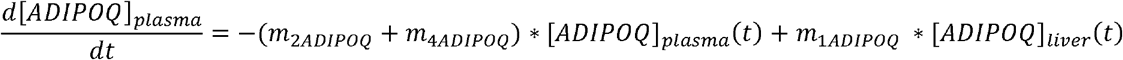

*m*_1*ADIPOQ*_ *m*_2*ADIPOQ*_ and *m*_3*ADIPOQ*_ (min^-1^) are rate parameters. *m*_4*ADIPOQ*_ is peripheral degradation. Hepatic extraction (HE) that determines liver degradation of adiponectin is described by:

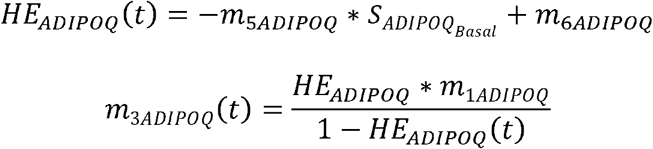

At basal steady state:

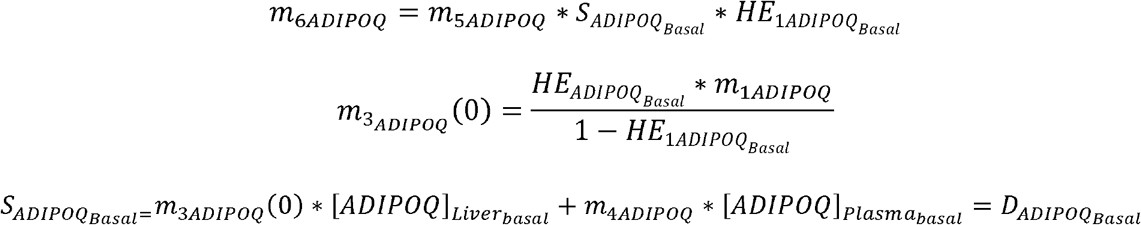

where *D_ADIPOQ_Basal__* is the basal total degradation.

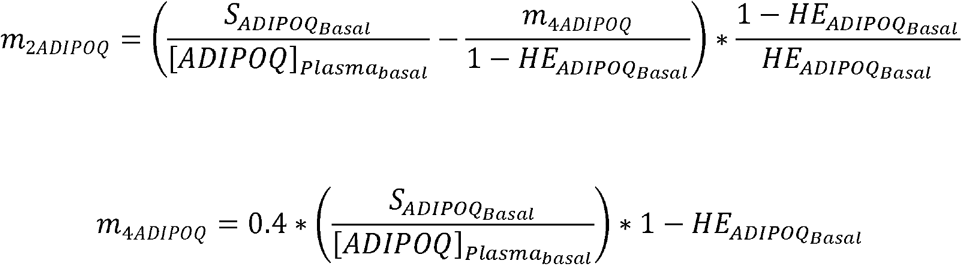

#### Hormone Concentration in Muscle and Interstitial Fluid

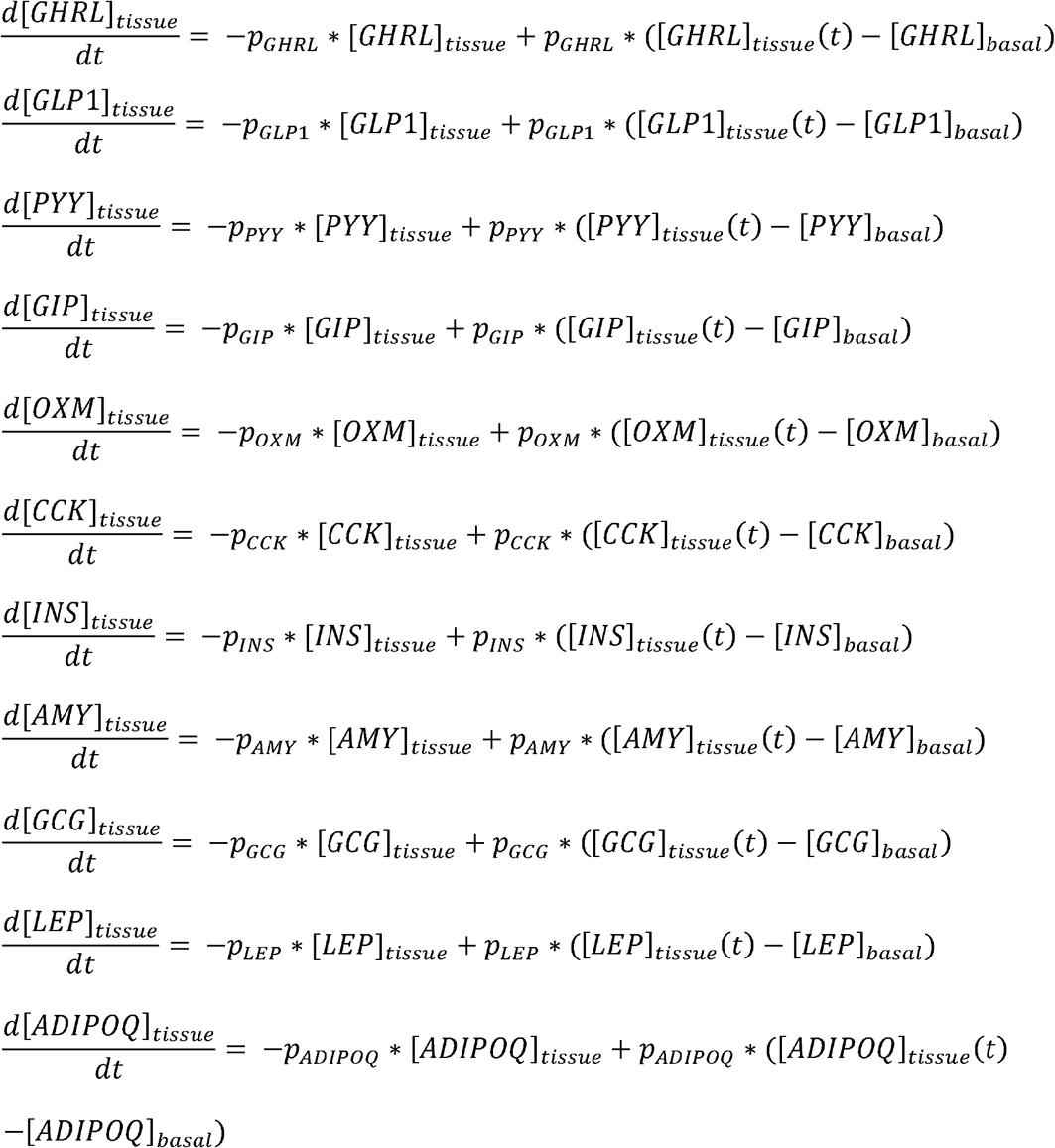

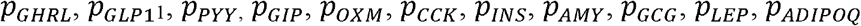 are the rate constants of the action on peripheral glucose utilization by ghrelin, GLP-1, PYY, GIP, oxyntomodulin, cholecystokinin, insulin, amylin, glucagon, leptin, adiponectin, respectively.

### Quantitative output of the model

The ratio of the Area Under the Curve (AUC) between postprandial state and fasting state was calculated for each hormone in the lean, obese and post-surgery state using MATLAB2018a trapezoidal function. (Fig 3B). These quantities utilized as outputs in the following parameter sensitivity analysis.

### Parameter sensitivity analysis (PSA) utilizing multivariable regression (MRA)

In order to assess the sensitivity of the kinetic parameters that regulate endocrinology in lean, obese and post-surgery subjects, we performed multivariable regression analysis [18].

The logic of the analysis is to randomly vary a set of inputs *n* times (trials), utilize the new set of inputs to generate an output according to the function in study and then perform a regression analysis to assess how much the variance in the parameter affects the output.

In our case, the kinetic parameters of the dynamical model were utilized as inputs. We varied the 141 parameters utilizing random scale factors from a log-normal distribution with a median value of 1. The distribution parameter σ, specifying the standard deviation of the distribution of log-transformed variables, controlled the extent to which parameters varied. We generated a total of 300 parameter sets that were stored in the input matrix, *I*(*300×141*). Each new set of parameters was utilized to simulate the dynamical model and the output (AUC ratios for each hormone) were stored in the output matrix, *O* (*300×11*).

Values in input and output matrices were mean-centered and normalized by standard deviations for each column. Since random-scale factors obeyed a log-normal distribution, most values in the input matrices were log-transformed before computing the standard deviation.

The PLS regression, performed on matrices I and O using the NIPALS algorithm [65, 66], produced a matrix of regression coefficients, *B_PLS_*(*141×11*). For each set of inputs, this matrix can be used to predict the resulting outputs. The regression determines *B_PLS_* such that *O_predicted_* = *I* x *B_PLS_* is close to the original output matrix *O*.

To test the susceptibility of our results to the number of trials, we performed the analysis for an increasing number of trials (500, 1000, 3000, 10000). PLS regression was performed using routines and methods from [67]. All computations were performed in MATLAB 2018a.

### Ranking of the kinetic parameters from PSA

The kinetic parameters were ranked according to their respective PLS coefficients [0–1]. The top ten most sensitive kinetic parameters for lean, obese and post-surgery subjects are shown in **Figure 4.**

### Association of kinetic parameters to their respective genes (MBCO)

To scale the previous results to a molecular level, the Molecular Biology of the Cell Ontology (MCBO) algorithm was used. Each parameter was used as a query in PubMed to retrieve the genes associated with it. To be able to identify these terms in our PubMed article sets, a dictionary that associates biological entities with key terms that might be used by authors when writing about the biological entity was generated. The MCBO algorithm screens the PubMed titles and abstracts of each SCP-specific article set and counts how many articles mentioned a certain gene at least once [20]. A minimum number of abstracts within an SCP specific abstract that is needed to mention a gene (level-1 SCPs: at least 4 abstracts, level-2 SCPs at least 3 abstracts, level-3 SCPs at least 2 abstracts, level-4 SCPs at least 1 abstract) is defined. Genes that were not mentioned in the specified number of abstracts were removed. These article counts for each gene-SCP association were subjected to statistical enrichment analysis to calculate the selectivity of each gene-SCP association, i.e. how selective is that particular gene for that particular SCP. Fisher’s exact test was used to calculate two different p-values, one with the genes of all SCPs of the same level (same level p-values), and one with the genes of all SCPs that have the same parent and therefore belong to the same children set (same children set p-values) as a background set.

### Average ranking of the genes utilizing PLS and MCBO score

The MCBO score assigned to each identified gene and the sensitivity rank of each kinetic parameter were utilized to calculate the average ranking for each identified gene.

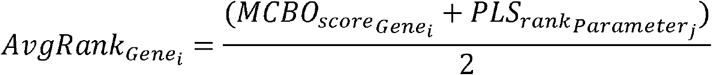

The MCBO score describes the significance of the association of a gene *i* to a specific parameter *j*. The PLS rank assesses the degree of sensitivity of that parameter and it is used to assess sensitivity variability between lean, obese and post-surgery subjects. The average ranking allows one to rank each gene according to its relevance in both the endocrinological control system and cellular processes.

### Weighting the unfiltered PPI network by nearest neighbor distance to an average ranked gene

The unfiltered PPI network created in the first step of the study was weighted according to the distance of each of its genes to the average ranked genes in lean, obese and post-surgery subjects. A nearest-neighbor algorithm was utilized to calculate the distances. The distance values (*0, 1,.,.n*) were utilized to weight the edges of the PPI network. This generated 3 physiologically specific PPI networks for lean, obese and post-surgery subjects, respectively.

### Peripheral tissue specific mRNA expression levels

The tissue expression of mRNAs of the genes in the lean, obese, and post-surgery subnetworks was retrieved from the GTEx Portal [19]. The GTEx_Analysis_v6p_RNA-seq_RNA-SeQCv1.1.8_gene_median_rpkm database (May 2017) was downloaded. The 75th percentile was used as a cut-off for high expression. Data for subcutaneous adipose tissue, visceral adipose tissue, stomach, pancreas, liver, intestine, and muscle were selected. The expression values used for adipose tissue were the means for subcutaneous and visceral tissues. Heatmaps of the expression levels for each physiologically-specific subnetwork are represented in **Figure 4B**.

### Tissue-specific PPI subnetworks for lean, obese and post-surgery subjects

Tissue-specific genes were utilized as seed genes to generate tissue-specific subnetworks for lean, obese and post-surgery subjects. We utilized the X2K software [5] to create a proteinprotein interaction (PPI) network. A path length of 2 was used, the other settings of the software are screenshot in (**Supplementary Figure S12 and S13**).

### Validation of tissue selective networks using drug targets from the FAERS database

We identified FDA-approved drugs that were reported in the FAERS database (June 2017) for inducing weight gain and weight loss [20]. A *p*-value <10^-5^ and number of reports>*1000* were used as a cut-off. The top 100 drugs causing weight gain and the top 100 drugs causing weight loss were so identified. Their respective targets were identified combining multiple databases: Drugbank [19], The Druggable Genome [68, 69], and Therapeutic Targets Database [70]. Targets that are exclusively expressed in the central nervous system were excluded. Targets which are expressed in adipose tissue, intestine, liver, muscle, pancreas, stomach but drugs acting on them do not cross the brain blood barrier were selected for the validation step. The number of tissue specific targets that were identifiable in the tissue-specific PPI subnetworks was used as validation of the reliability of the previous steps of the study.

### Catabolic and anabolic subcellular processes

From the previous analysis, we calculated the number of significant catabolic and anabolic processes and the ratio of significant catabolic and anabolic processes in lean, obese, and post-surgery subjects (Figure 6A). The tissue specific ratio of significant catabolic and anabolic processes in lean, obese, and post-surgery is shown in (Figure 6B).

## Acknowledgments

This study was supported by a research grant from GSK to ISMMS and Systems Biology Center Grant P50GM071558.

## Author contributions

RI, DKR, V.B. and CVH. conceived and planned the study. IT and VB designed the computational framework and analyzed the genomic, endocrine and network data. CVH performed the FAERS data analysis. RI, PS and DKR supervised the project. EAS supervised the dynamical modeling studies. I.T, VB, EAS, JG, DKR and RI contributed to the interpretation of the results. IT, VB, JG and RI took the lead in writing the manuscript. All authors provided critical feedback and helped shape the research, analysis, and manuscript.

