## Supplementary material for "A tissue- and organ-based cell biological atlas of obesity-related human genes and cellular pathways": Supplementary Figure S9.pdf

**Multivariable Regression Analysis (MRA)**

Baseline kinetic parameters are randomly varied *n* times (simulated individuals)

↓

**Input matrix**
*300 simulated individuals* x *141 parameters*

↓

Run simulations for each new set of parameters

↓

**Output matrix**
*300 simulated individuals* each with *11 AUC ratios*

↓

**Partial Least Square (PLS) MRA**

Changes in the outputs can be approximated as the change in parameters (input) times a matrix of parameter sensitivities, PLS coefficient matrix (*141 x 11*) (*center*)

↓

**Kinetic parameters ranked according to the PLS coefficients**

↓

**Identification of top 10 sensitive kinetic parameters**

*Repeat for 300, 500, 1000, 3000, 10000 simulated individuals*

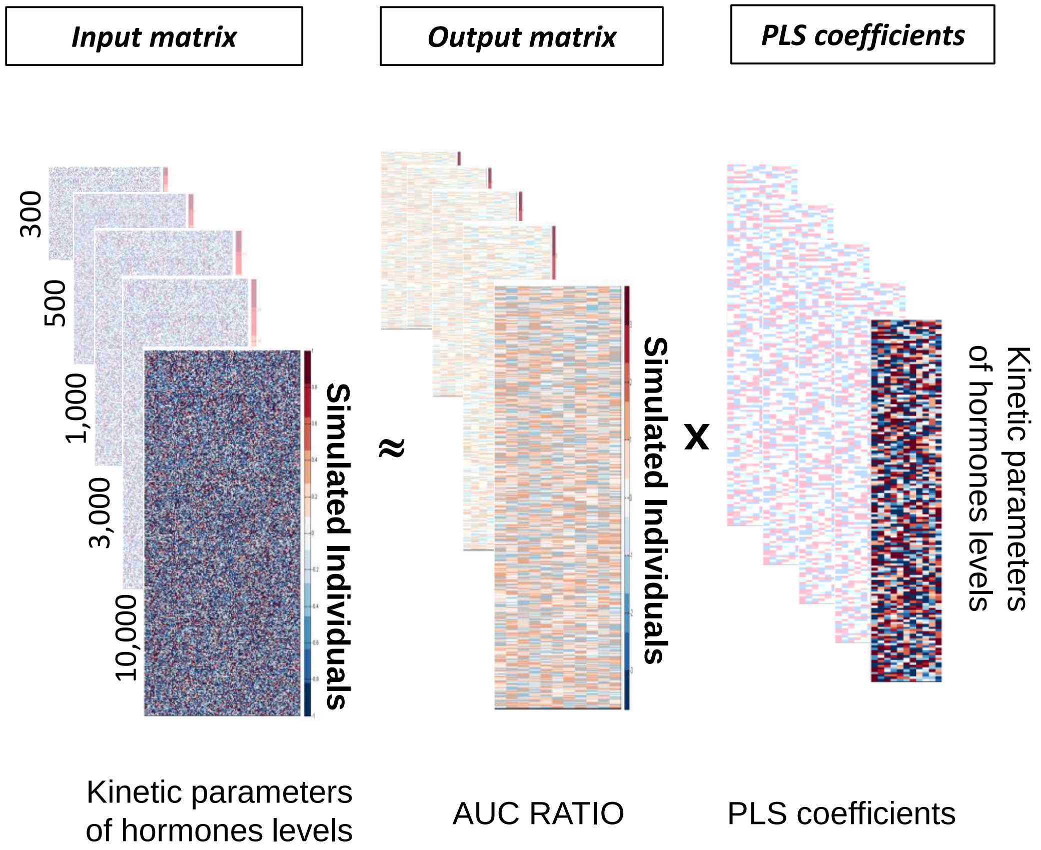
