## Supplementary material for "A tissue- and organ-based cell biological atlas of obesity-related human genes and cellular pathways": Supplementary Table S1.pptx

### Slide 1
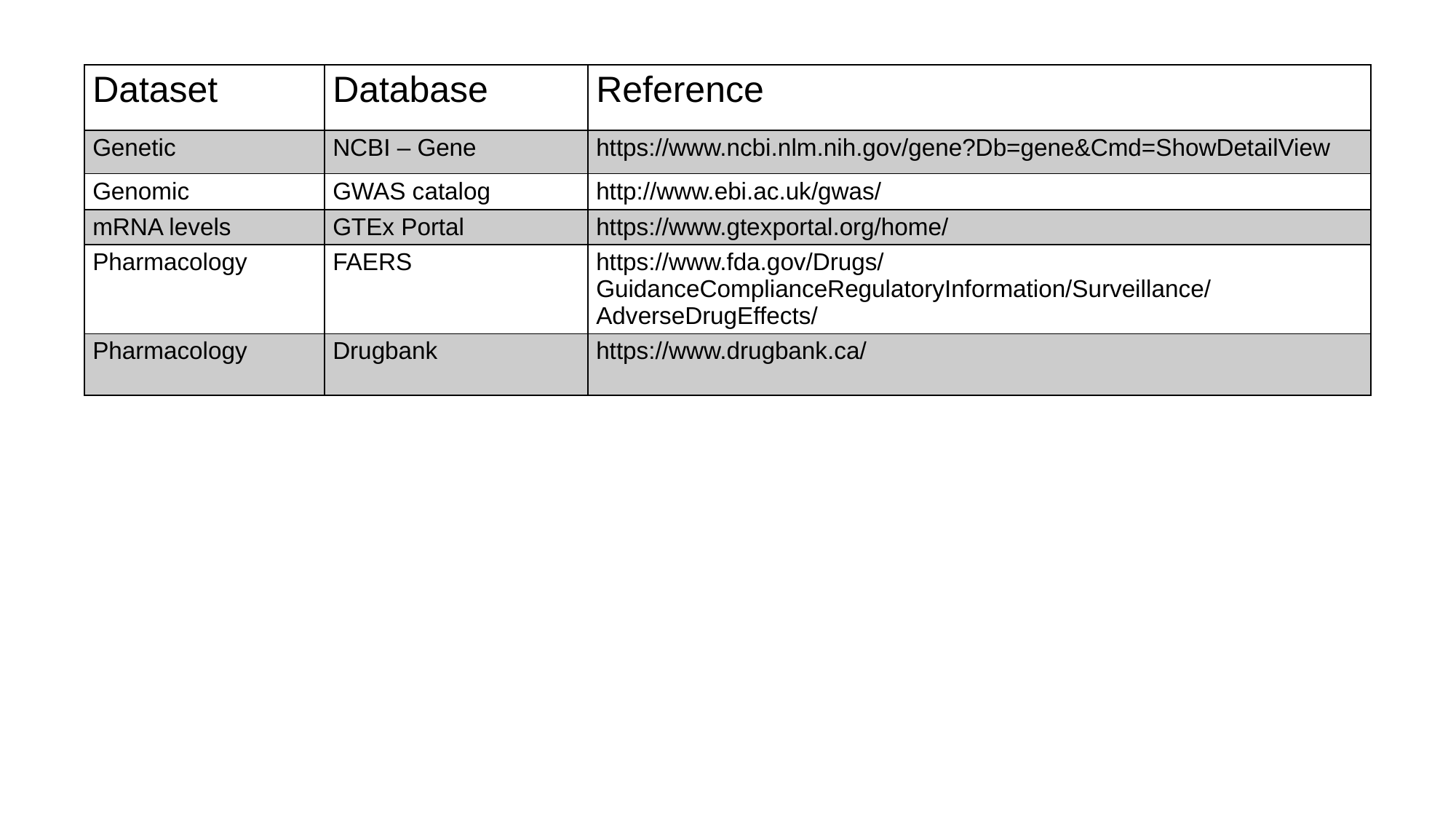

| Dataset | Database | Reference |
| --- | --- | --- |
| Genetic | NCBI – Gene | https://www.ncbi.nlm.nih.gov/gene?Db=gene&Cmd=ShowDetailView |
| Genomic | GWAS catalog | http://www.ebi.ac.uk/gwas/ |
| mRNA levels | GTEx Portal | https://www.gtexportal.org/home/ |
| Pharmacology | FAERS | https://www.fda.gov/Drugs/GuidanceComplianceRegulatoryInformation/Surveillance/AdverseDrugEffects/ |
| Pharmacology | Drugbank | https://www.drugbank.ca/ |
