## Supplementary material for "A tissue- and organ-based cell biological atlas of obesity-related human genes and cellular pathways": Supplementary Table S2.pptx

### Slide 1
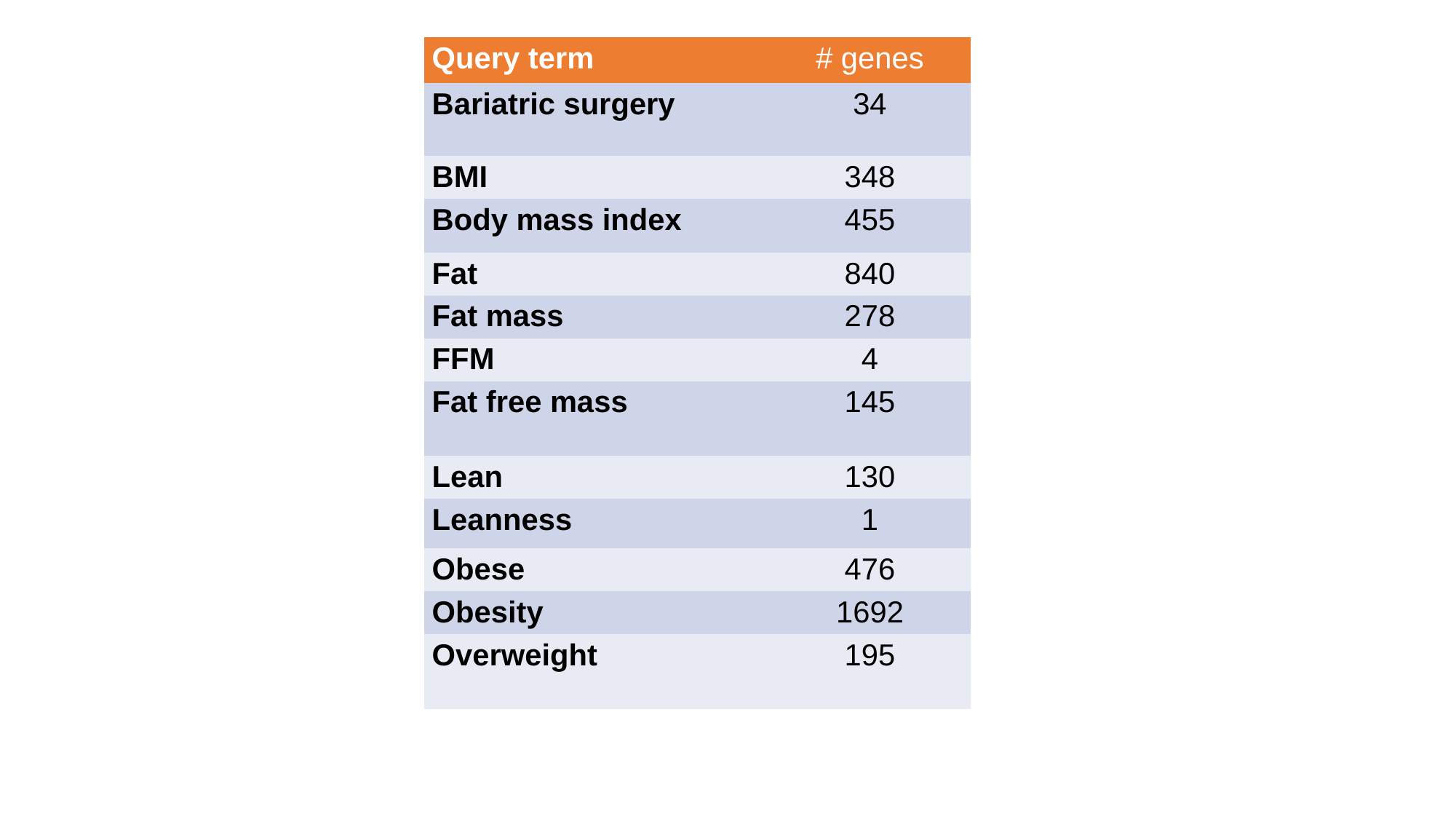

| Query term | # genes |
| --- | --- |
| Bariatric surgery | 34 |
| BMI | 348 |
| Body mass index | 455 |
| Fat | 840 |
| Fat mass | 278 |
| FFM | 4 |
| Fat free mass | 145 |
| Lean | 130 |
| Leanness | 1 |
| Obese | 476 |
| Obesity | 1692 |
| Overweight | 195 |
