## Supplementary material for "A tissue- and organ-based cell biological atlas of obesity-related human genes and cellular pathways": Supplementary Table S3.pptx

### Slide 1
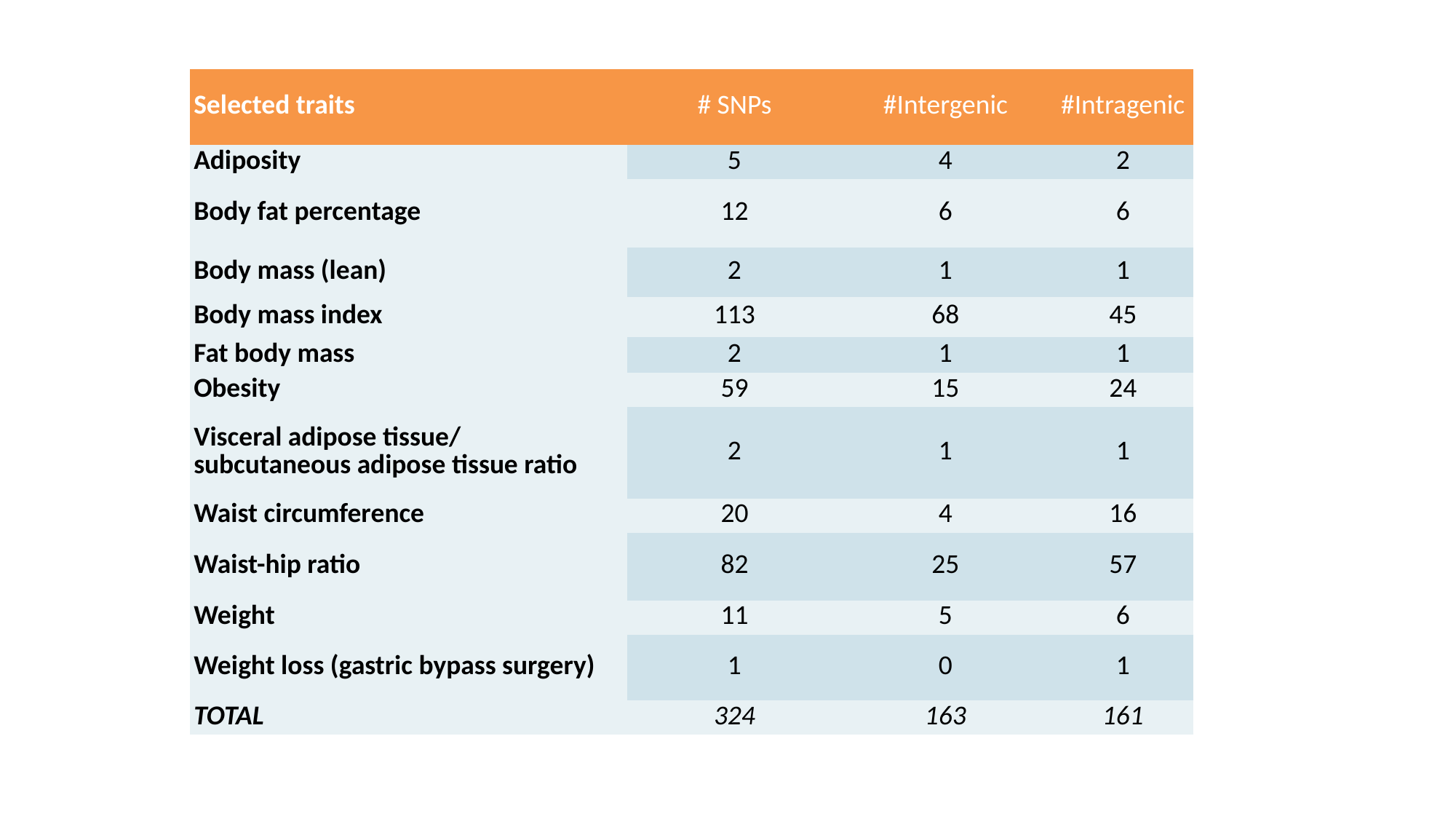

| Selected traits | # SNPs | #Intergenic | #Intragenic |
| --- | --- | --- | --- |
| Adiposity | 5 | 4 | 2 |
| Body fat percentage | 12 | 6 | 6 |
| Body mass (lean) | 2 | 1 | 1 |
| Body mass index | 113 | 68 | 45 |
| Fat body mass | 2 | 1 | 1 |
| Obesity | 59 | 15 | 24 |
| Visceral adipose tissue/ subcutaneous adipose tissue ratio | 2 | 1 | 1 |
| Waist circumference | 20 | 4 | 16 |
| Waist-hip ratio | 82 | 25 | 57 |
| Weight | 11 | 5 | 6 |
| Weight loss (gastric bypass surgery) | 1 | 0 | 1 |
| TOTAL | 324 | 163 | 161 |
