## Supplementary figures and images for "A tissue- and organ-based cell biological atlas of obesity-related human genes and cellular pathways"

### Supplementary Figure S1.pdf

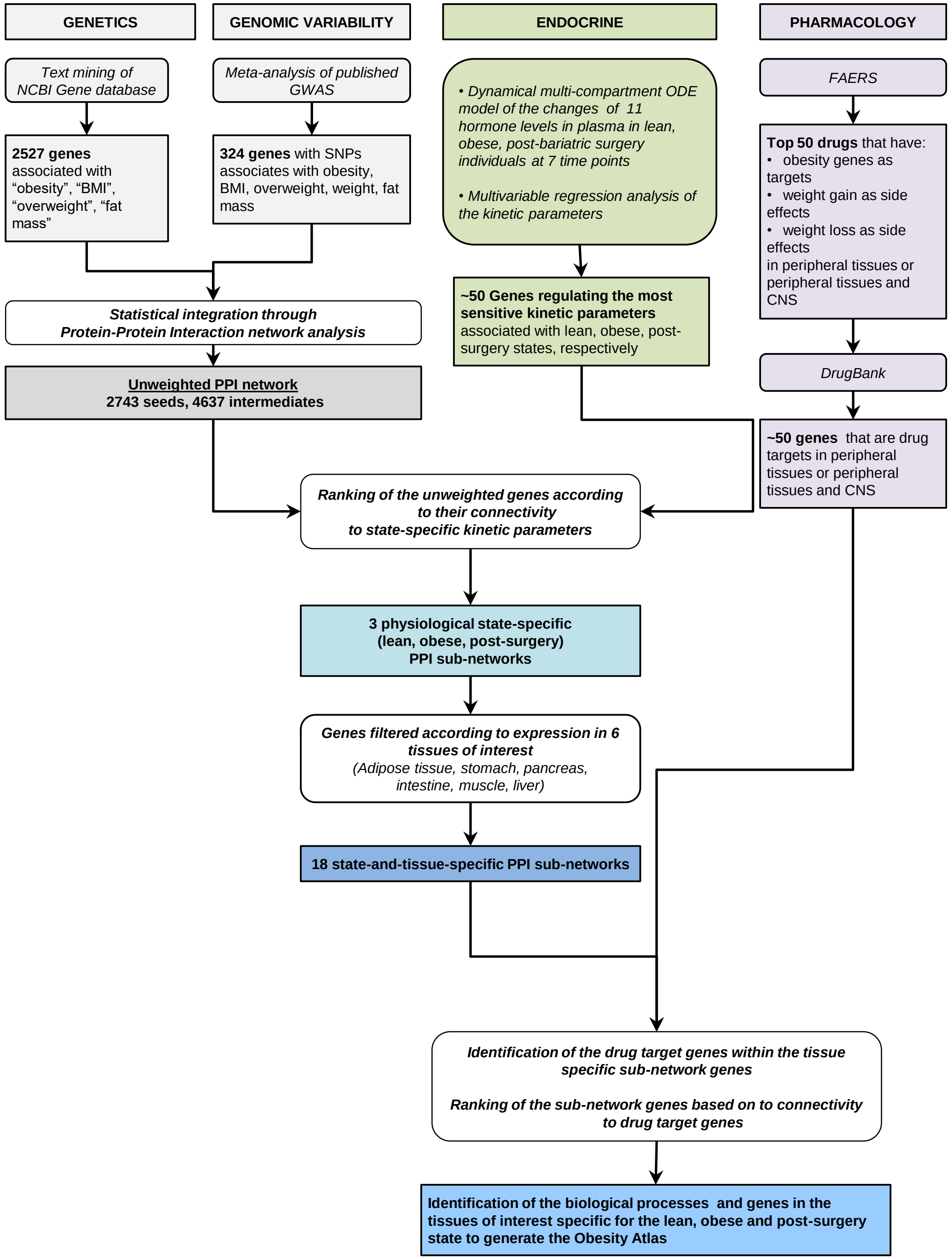

### Supplementary Figure S2.pdf

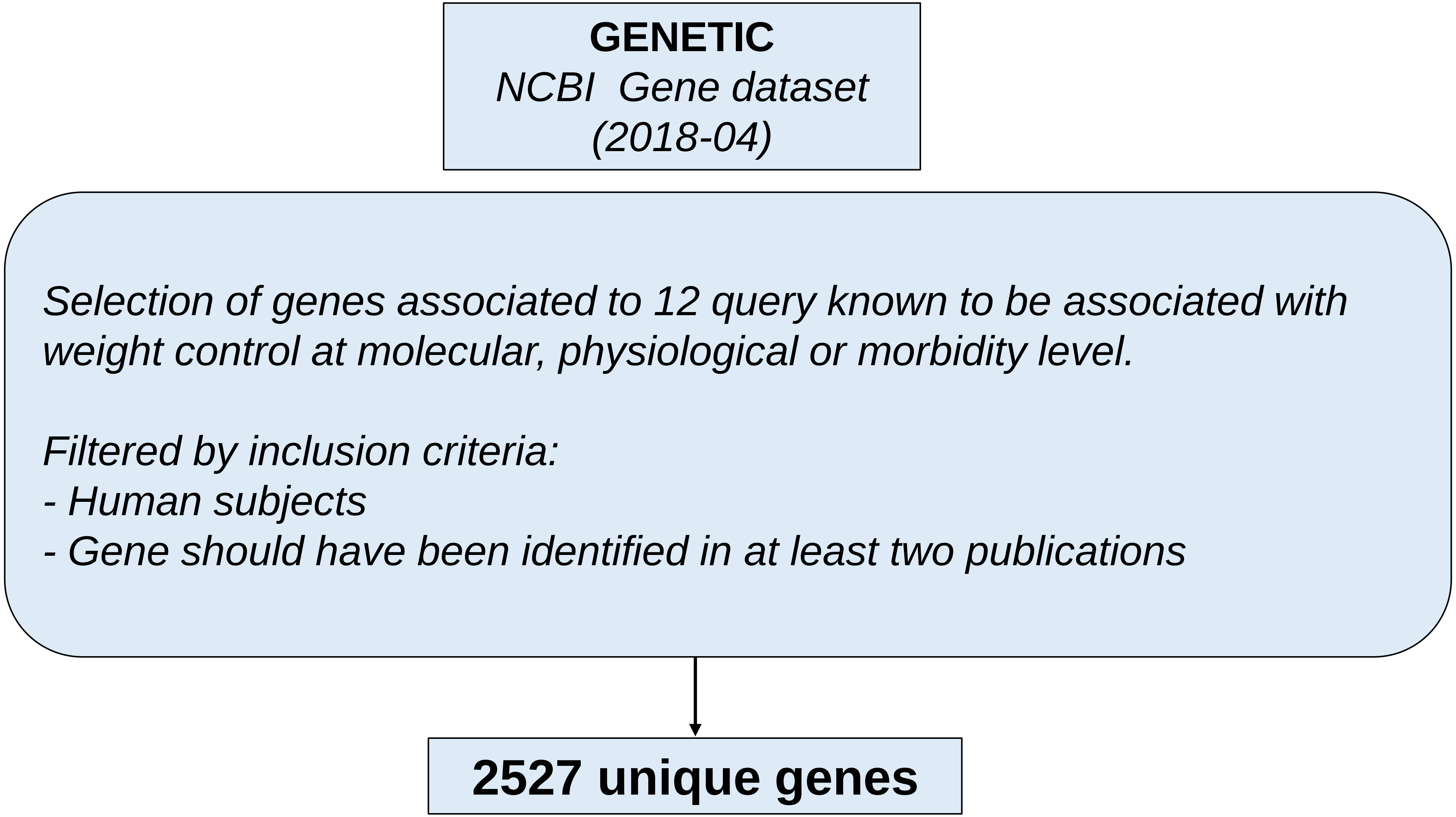

### Supplementary Figure S3.pdf

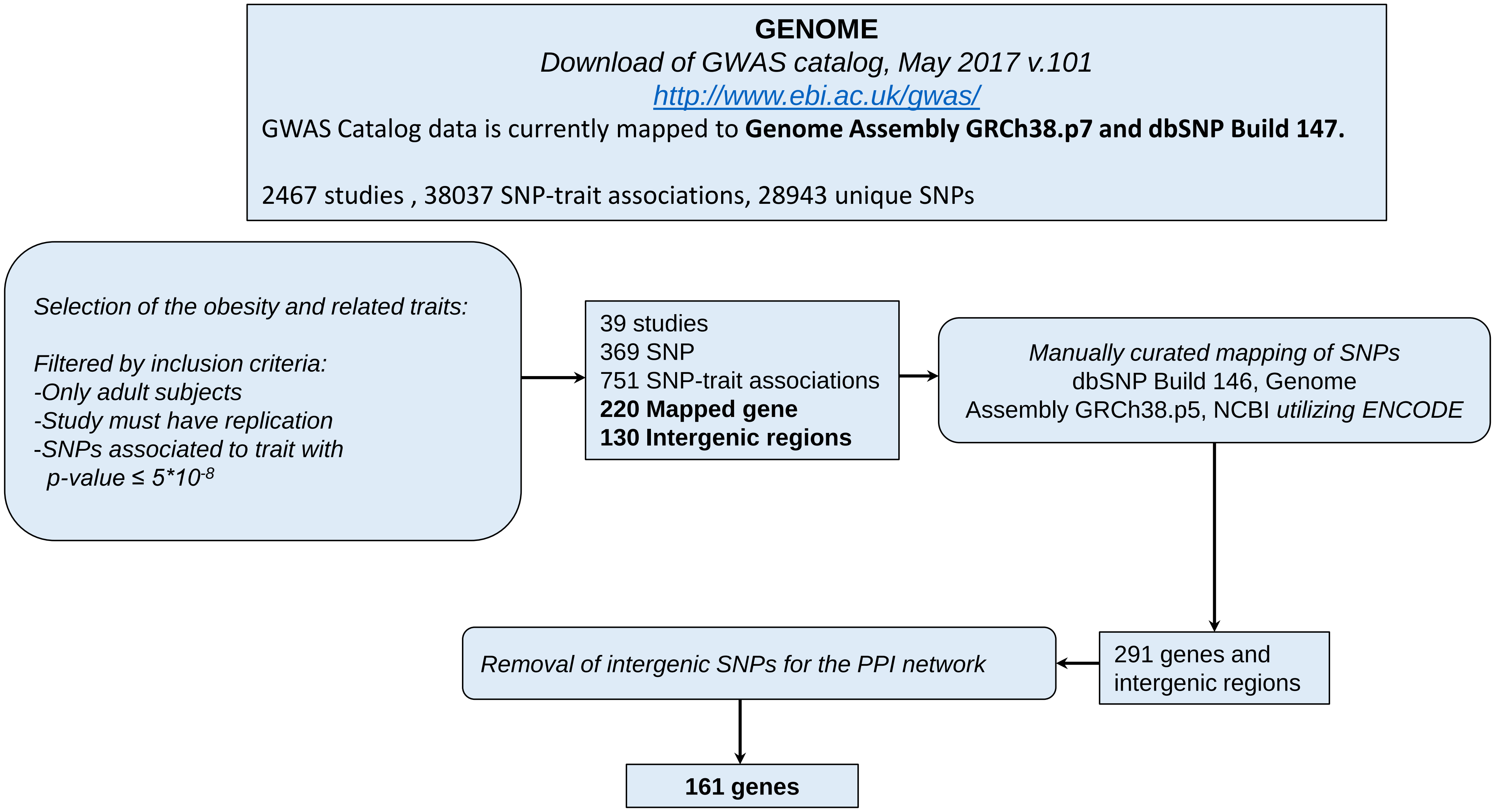

### Supplementary Figure S4.pdf

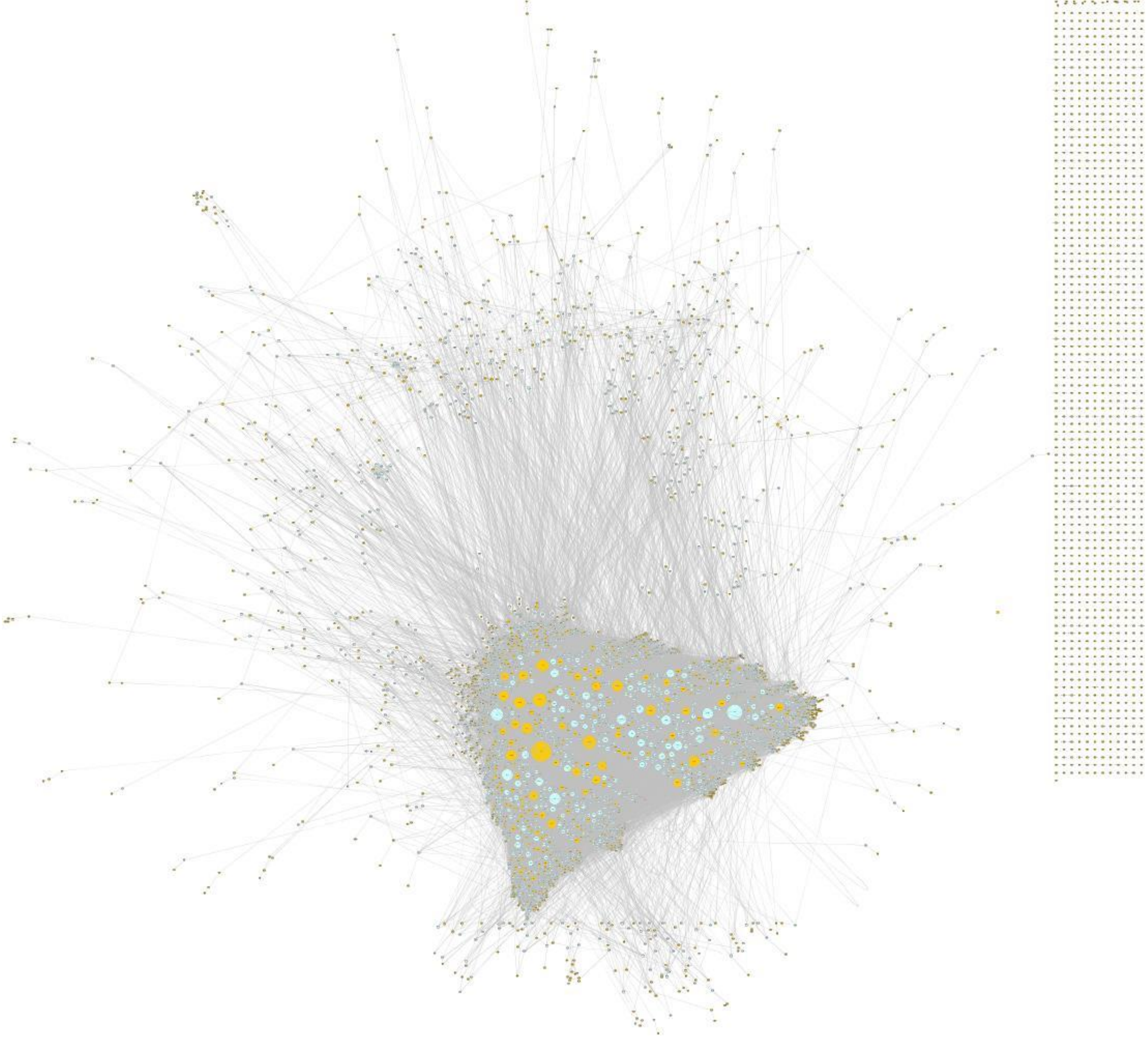

### Supplementary Figure S5.tif

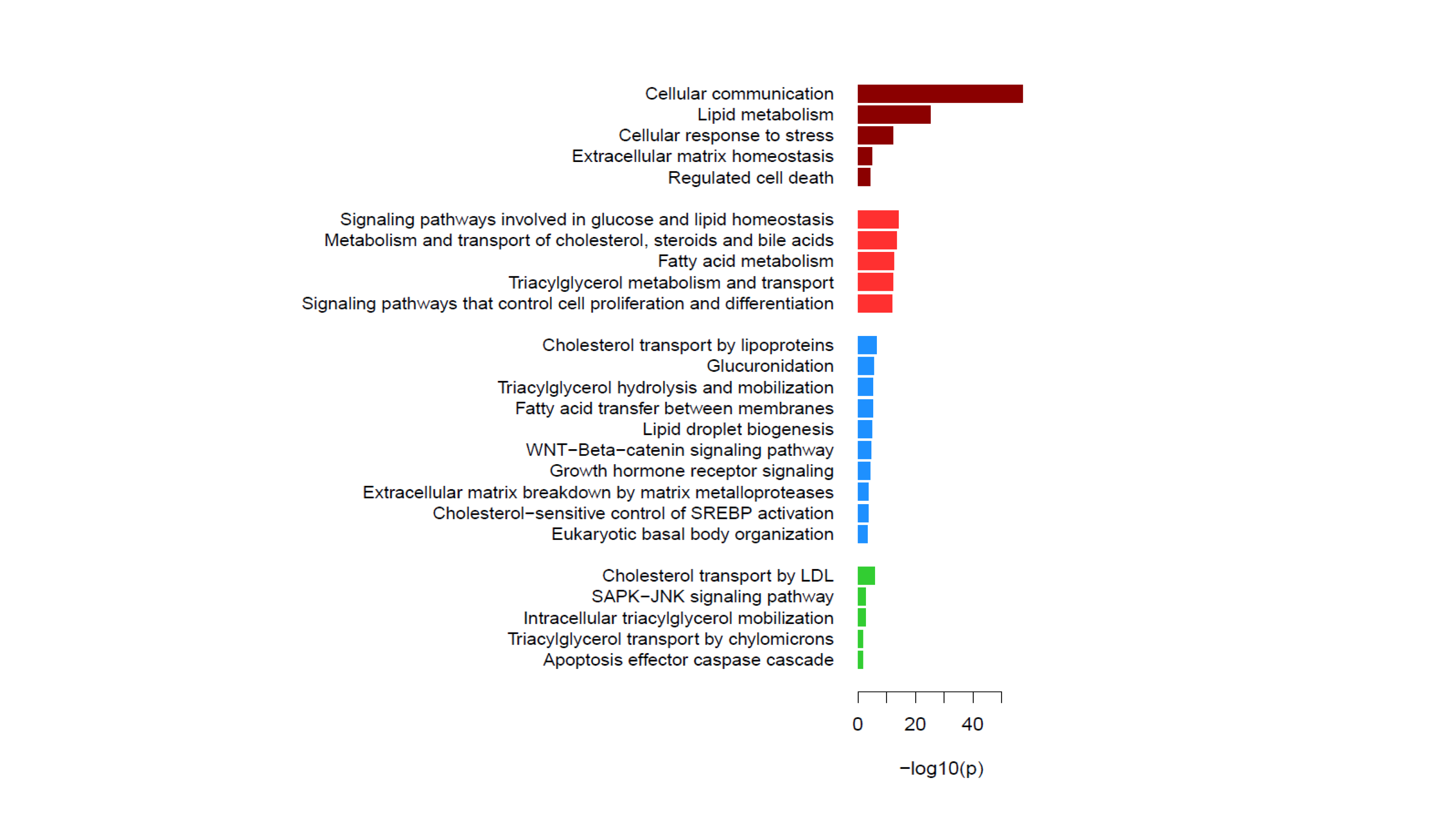

### Supplementary Figure S6.tif

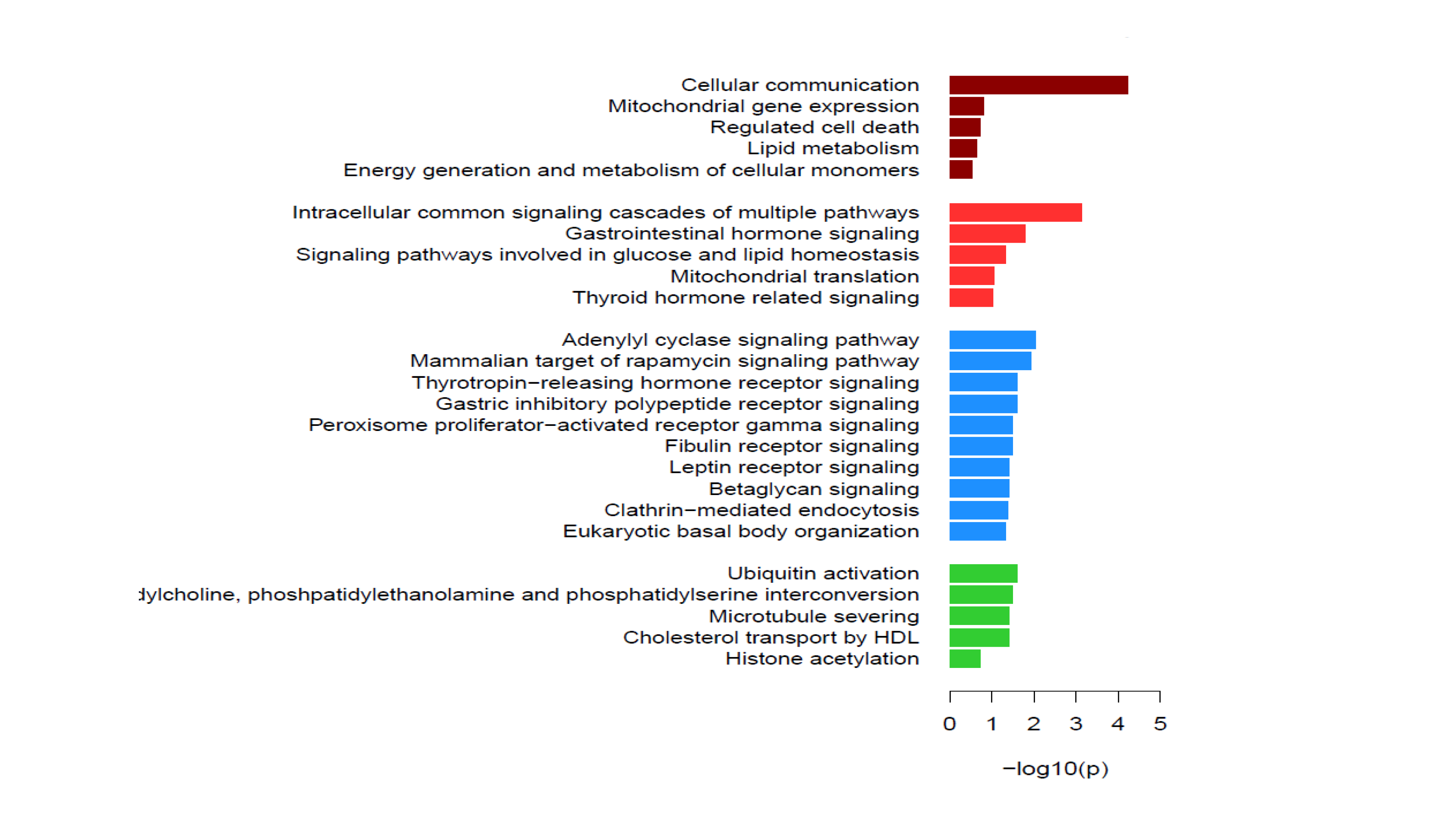

### Supplementary Figure S7.tif

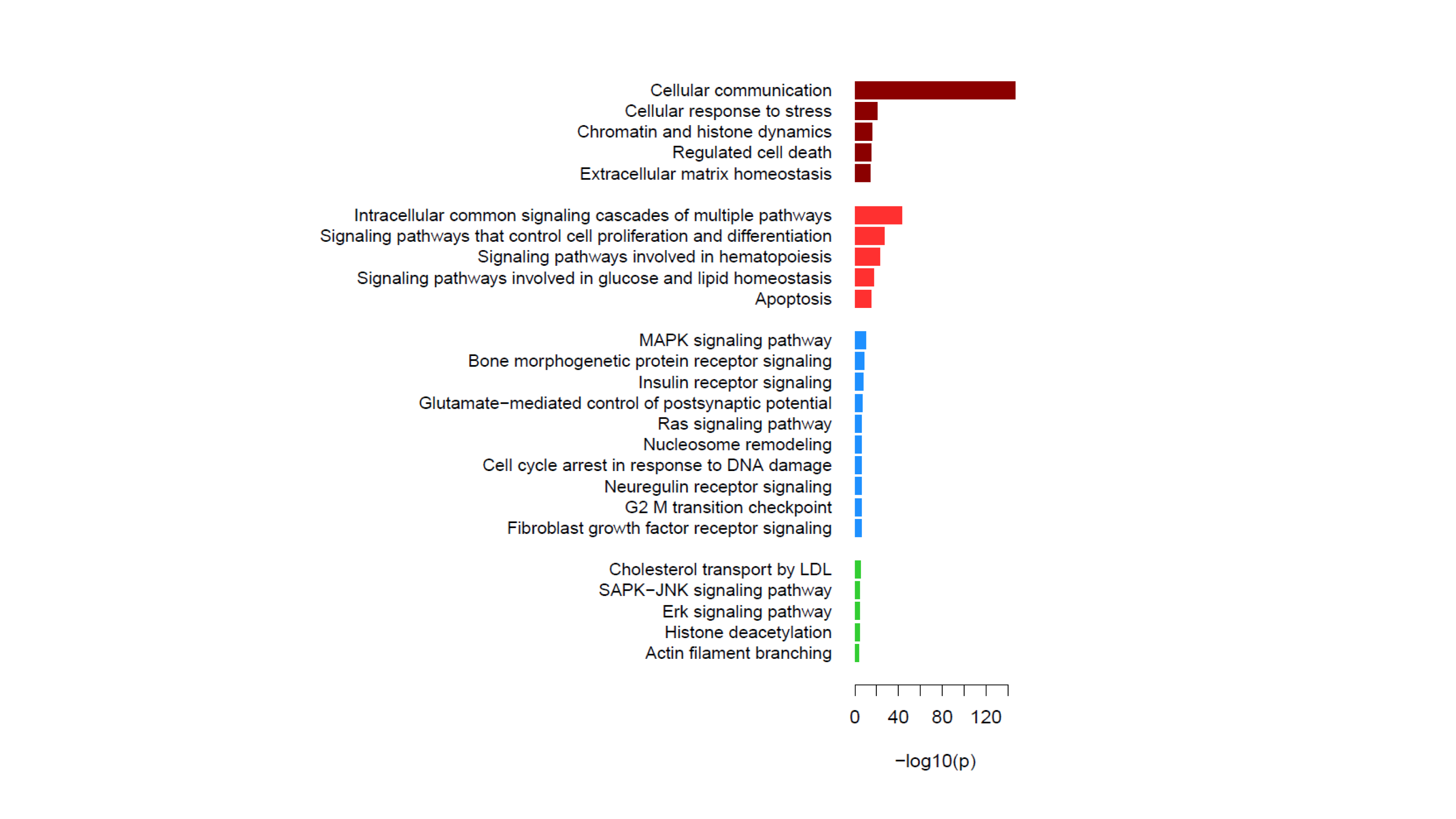

### Supplementary Figure S8.pdf

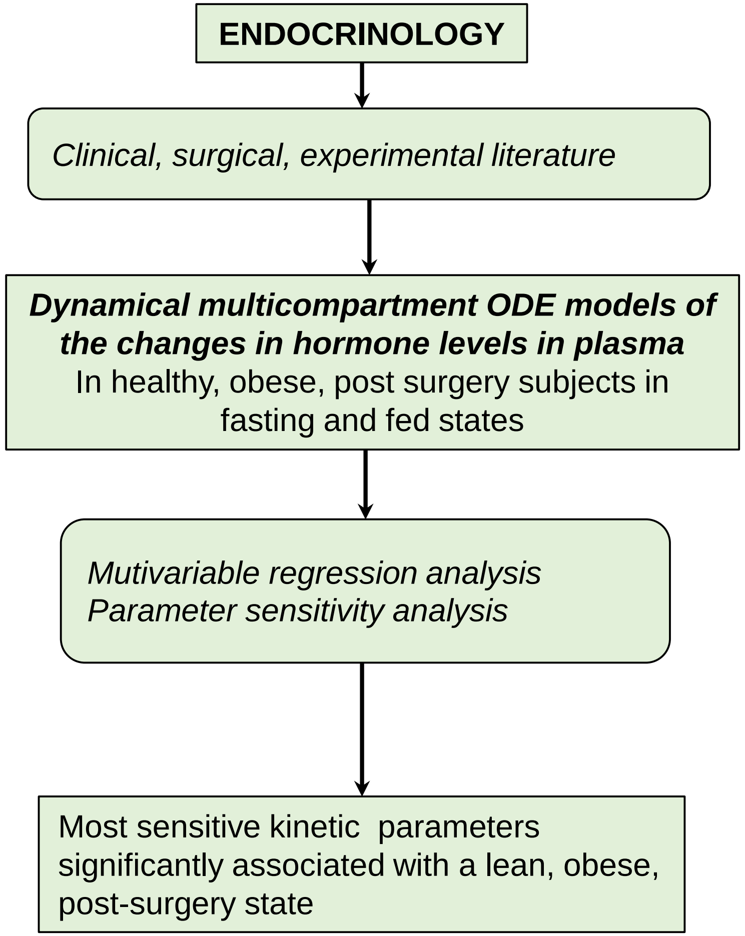

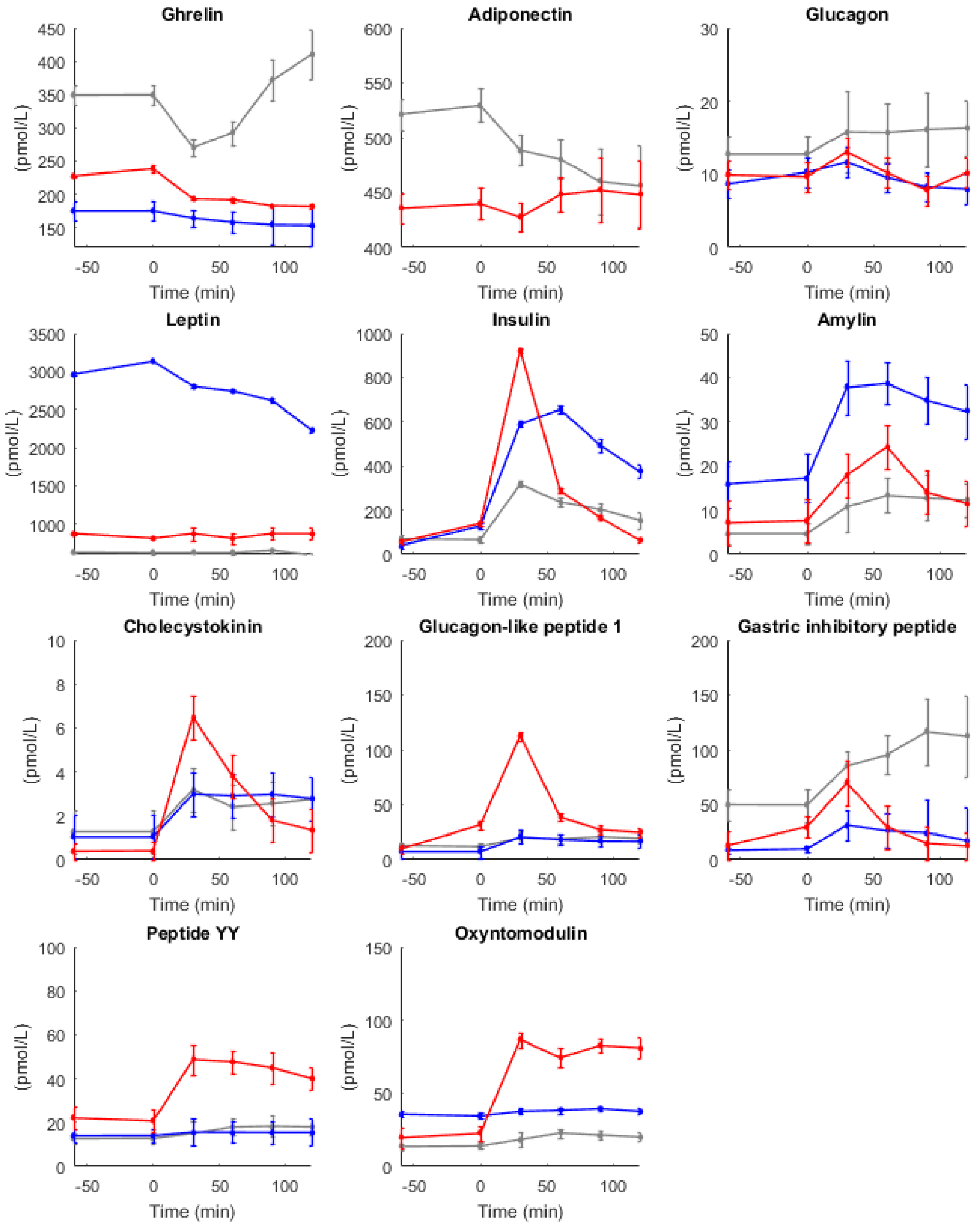

### Supplementary Figure S10.pdf

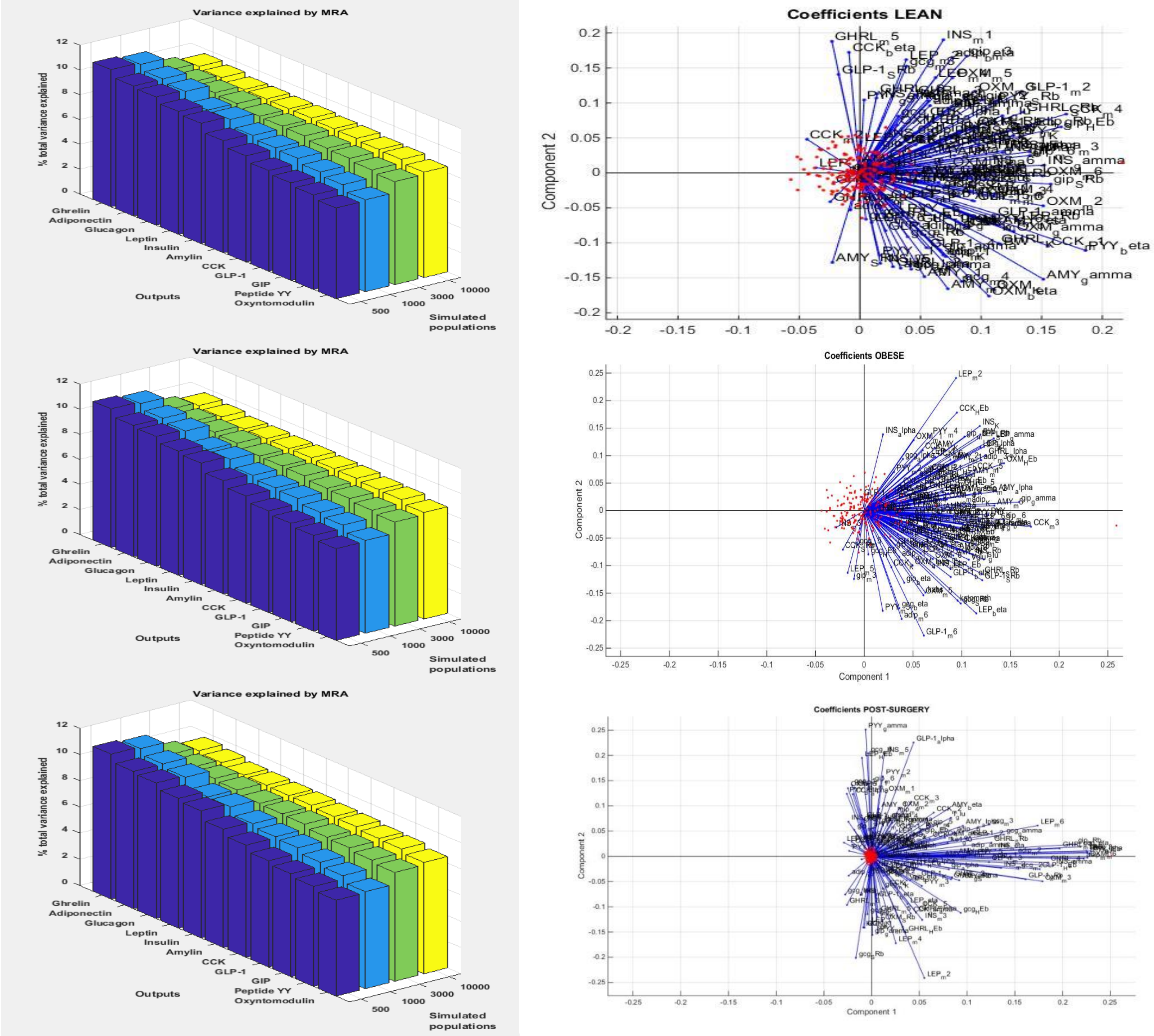

### Supplementary Figure S11.pdf

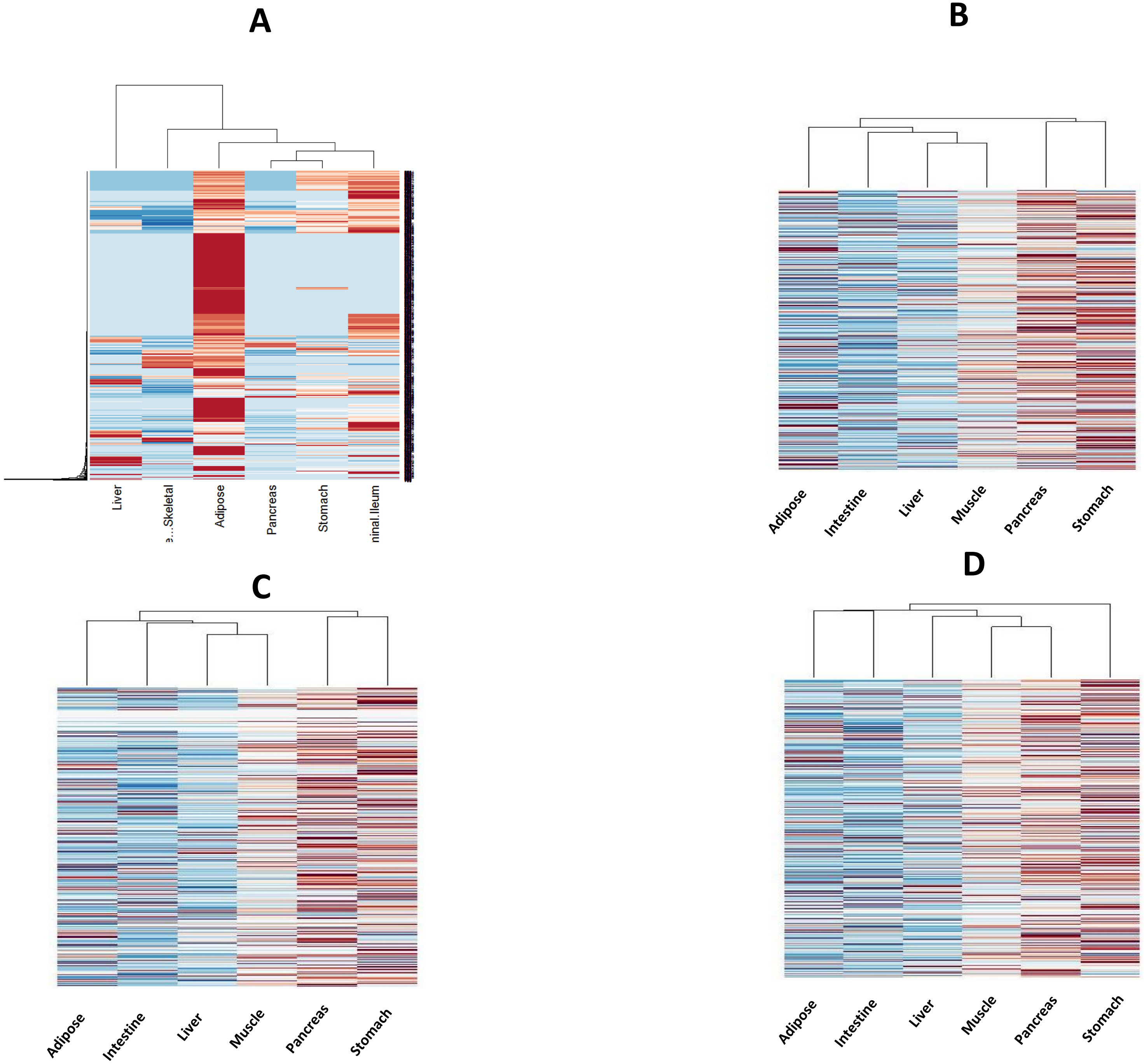

### Supplementary Figure S12 .pdf

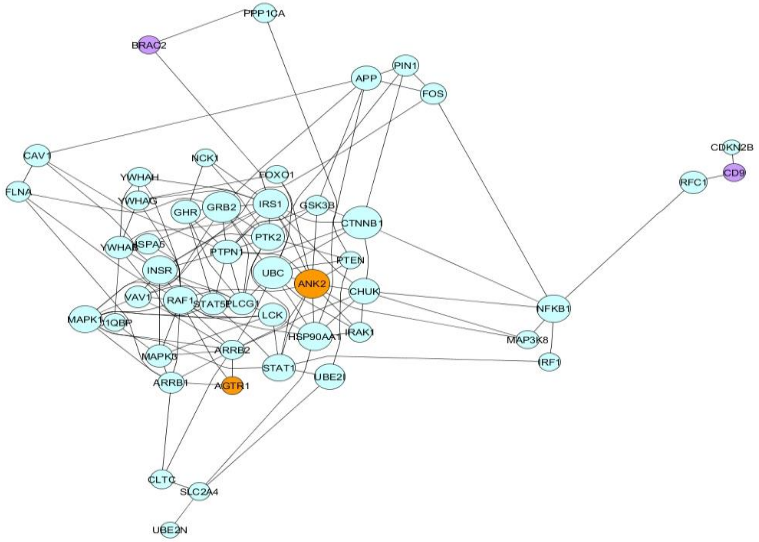


**Adipose tissue**

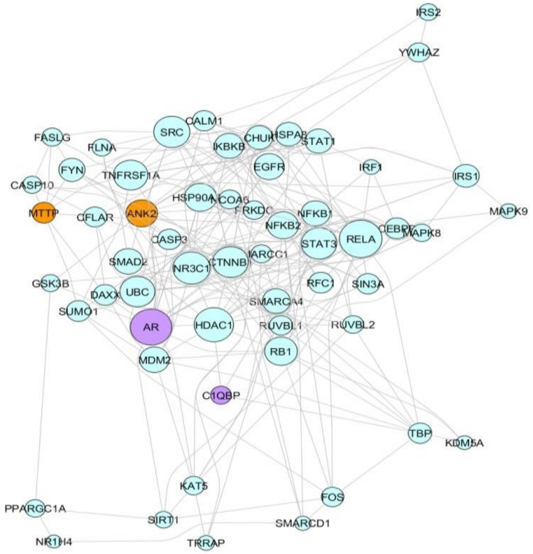


**Intestine**

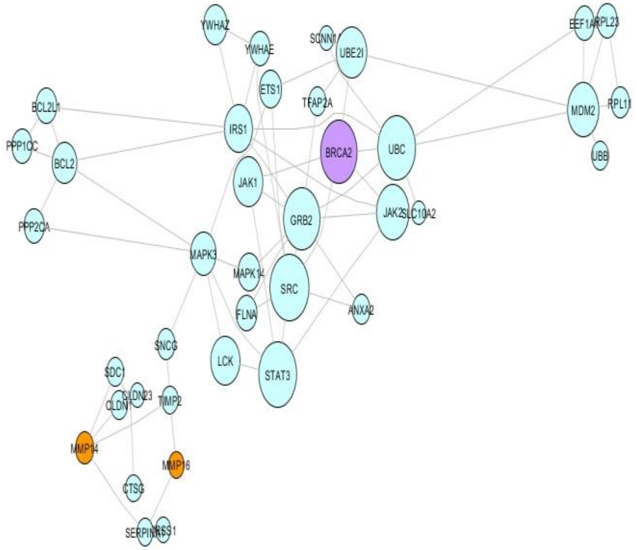


**Liver**

### Supplementary Figure S13.pdf

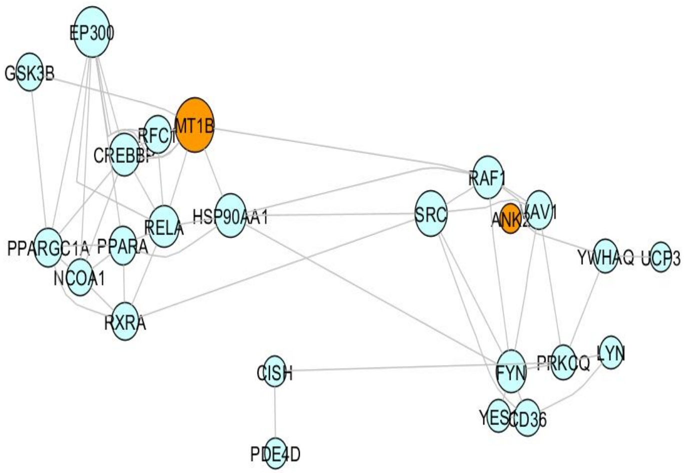


**Muscle**

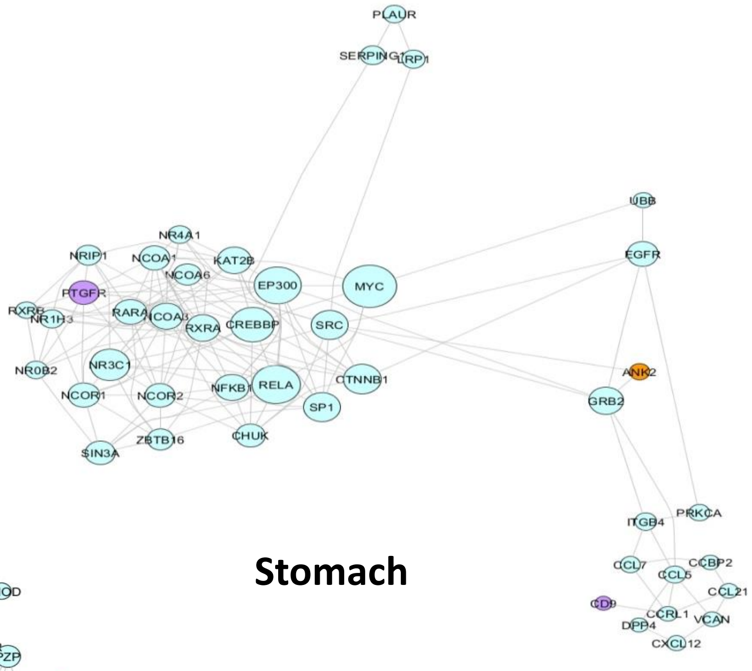


**Stomach**

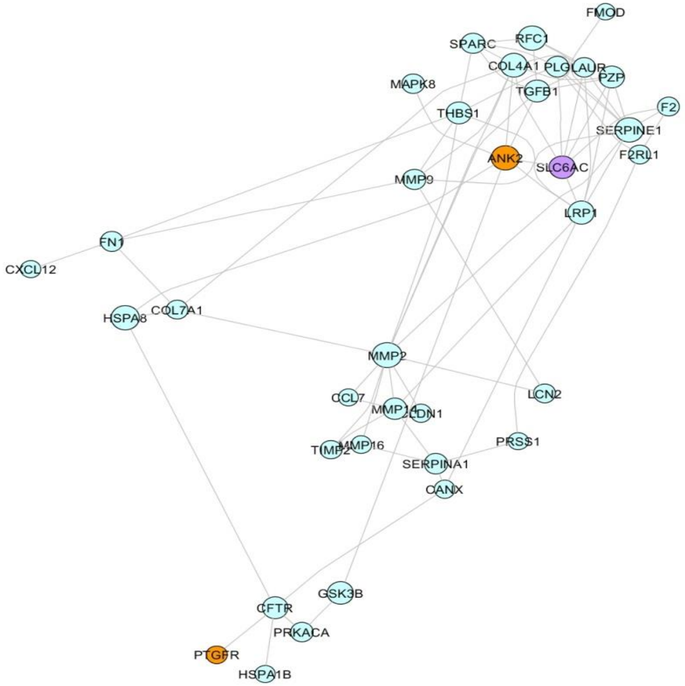


**Pancreas**

### Supplementary Figure S14.pdf

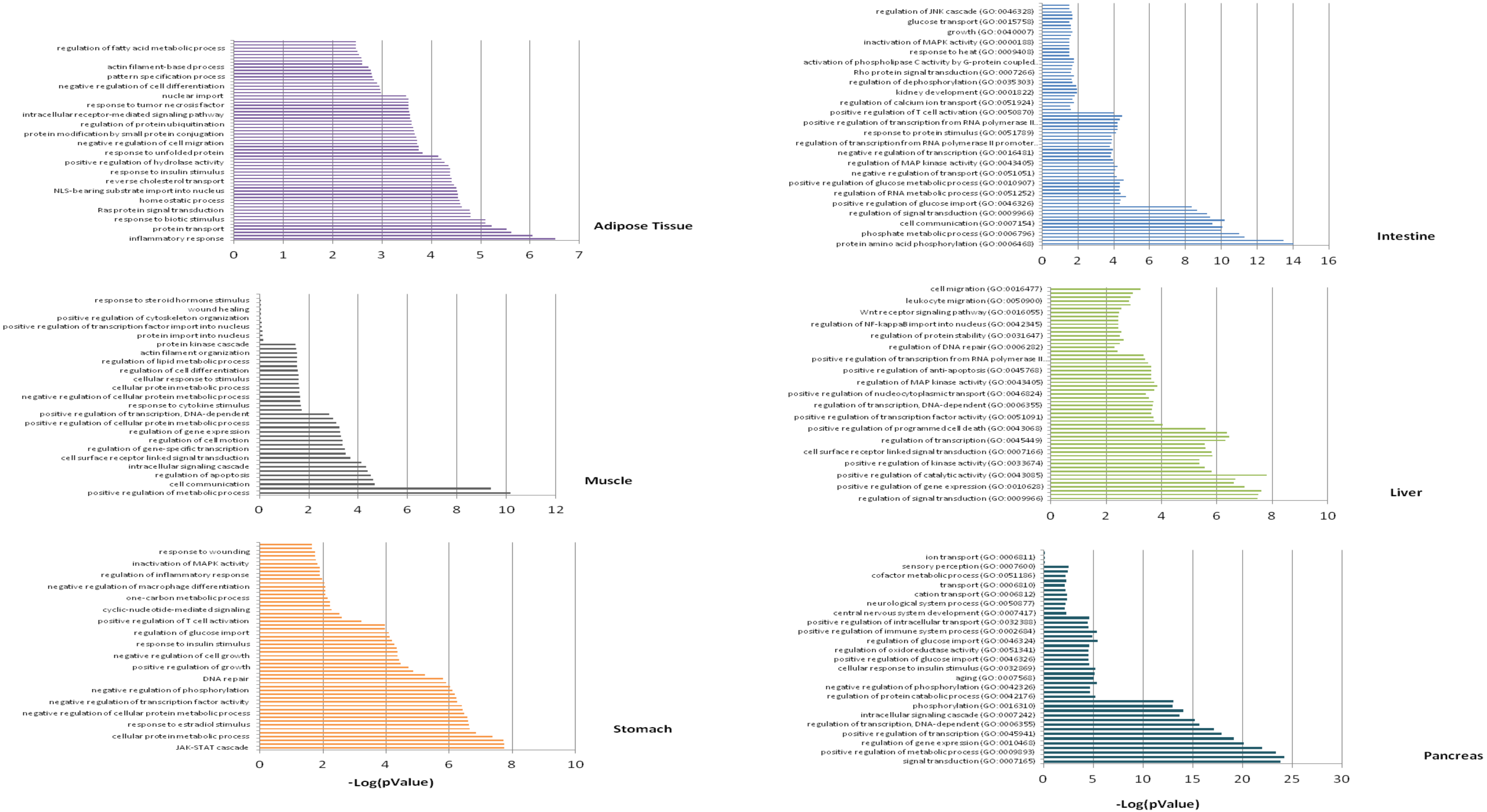
